# Senescence-Linked Fibrosis in the Aging Human Ovary Revealed by p16-Based Histological Profiling and Spatial Transcriptomics

**DOI:** 10.64898/2025.12.03.692228

**Authors:** Mark A. Watson, Pooja Raj Devrukhkar, Natalia F. Murad, Fan Wu, Moo Joong Kim, Hannah Anvari, Uyen Tran, Nicholas Martin, Tommy Tran, Giuliana Zaza, Kevin Schneider, Bikem Soygur, Elisheva D. Shanes, Denis Wirtz, Mary Ellen G. Pavone, Simon Melov, Pei-Hsun Wu, David Furman, Francesca E. Duncan, Birgit Schilling

**Author notes:** Co-correspondence: Birgit Schilling, Ph.D. Buck Institute for Research on Aging, Novato, CA 94945, USA. Francesca E. Duncan, Ph.D. Department of Obstetrics and Gynecology, Feinberg School of Medicine, Northwestern University, Chicago, IL, 60611 USA.

## Abstract

Cellular senescence is implicated as a driver of ovarian aging, but senescent cells in the human postmenopausal ovary remain poorly defined. Using spatially resolved p16^INK4a^ protein expression, a canonical senescence marker, we identified and mapped senescent cells in postmenopausal ovaries. We integrated p16 immunohistochemistry, multiplexed immunofluorescence, spatial transcriptomics, and AI-guided digital pathology to map senescent microenvironments. p16-positive cells formed discrete stromal, vascular, and cyst-associated clusters that increased with age and were enriched for macrophages and myofibroblast-like cells. Whole-transcriptome profiling of 92 spatial regions uncovered a 32-gene p16-associated signature, BuckSenOvary, that distinguished p16-positive regions across cortex and medulla. BuckSenOvary is characterized by suppression of cell-cycle regulators and activation of inflammatory and extracellular-matrix remodelling genes. AI-based collagen matrix analysis confirmed that p16-positive regions exhibit more architecturally complex collagen, demonstrating that focal senescent microenvironments are fibro-inflammatory. These findings position senescent ovarian niches as therapeutic targets to preserve ovarian function.

## Main

While women globally live ∼5 years longer than men, they experience a ∼2.4-year greater gap between lifespan and healthspan than men, spending a greater proportion of late life in poorer health^1,2^. The factors that contribute to this disparity are complex, but the impact of menopause on women’s health and aging is substantial. Menopause, defined as the permanent cessation of menstruation for at least 12 consecutive months, typically occurs around age 50 and reflects the cessation of ovarian function^3^. Beyond loss of fertility, menopause is associated with an abrupt decline in ovarian hormones, which increases systemic health risks, including cardiovascular disease, osteoporosis, and cognitive decline^4,5^. In fact, women who undergo natural menopause later generally live longer and have reduced risks of cardiovascular disease and all-cause mortality^6–8^. Due to medical and health advances, women may spend more than one-third of their lives in a postmenopausal state and experience the negative health sequelae. Thus, understanding the drivers of ovarian aging and how the post-menopausal ovary, long assumed to be inert, may contribute to systemic aging and disease is an emerging priority.

The human ovary is among the first organs to undergo functional decline with age, reflected not only by depletion and reduced quality of the oocyte pool but also by pronounced remodeling of the surrounding stromal microenvironment^9,10^. The aging ovarian microenvironment is characterized by increased fibrosis, changes in extracellular matrix composition, decreased vascularity, immune cell infiltration, and increased tissue stiffness^11–14^. Emerging technologies are revealing the transcriptomic landscape of the aging ovary^15–21^. However, the molecular and tissue-level drivers of postmenopausal ovarian fibro-inflammaging remain poorly defined.

Among the cellular mechanisms and hallmarks of aging^22^, cellular senescence is a key driver of age-related tissue dysfunction, promoting chronic inflammation and fibrosis through the senescence-associated secretory phenotype (SASP)^23–30^. Such secretomes foster tissue degradation, fibrotic remodeling, and immune evasion^23,31–33, 34^. Senescent cell accumulation is linked to aging in multiple organs, including skin , lung^35^, liver^36^, and kidney^37^, suggesting it may play similar roles in the aging ovary. While ovarian transcriptomic studies indicate age-associated senescence, and markers such as p16^INK4a^ (CDKN2A) and p21^CIP1^ (CDKN1A) increase with age, the spatial positioning and histological features of these cells in the postmenopausal human ovary remain unclear^16–18,20,21,38,39^. The heterogeneity of senescent cells across tissues, combined with the ovary’s diverse cellular composition and compartments, makes spatial and phenotypic resolution critical for gaining functional insight.

Here, we aimed to identify an ovarian senescence signature associated with aging and to characterize the niche of senescent-like cells and their effects within ovarian tissue. To this end, we developed a targeted spatial-molecular approach to identify and map senescent cells in native postmenopausal human ovaries using p16INK4a (p16) (Fig. 1a). We chose p16 as our primary marker because it is the most widely recognized canonical senescence-associated marker in both experimental models and human tissues^40–42^. p16, encoded by CDKN2A, is a cyclin-dependent kinase inhibitor that blocks CDK4/6 to enforce retinoblastoma (RB)-mediated G1 cell-cycle arrest^43–45^. Its expression increases in aging tissues, and sustained p16 indicates cells that have undergone replicative or stress-induced damage and have entered a stable growth arrest^41,46–48^. Distinct from cancerous ovarian tissue, where p16 expression exhibits intense block positivity staining, in this study, the analysis of 45 human post-menopausal healthy ovaries revealed sporadic, discrete clusters of p16positive cells throughout the stroma. The p16-positive signal occupied roughly 0.03-2.8% of tissue sections with a tendency towards increased expression with age.

**Figure 1.**
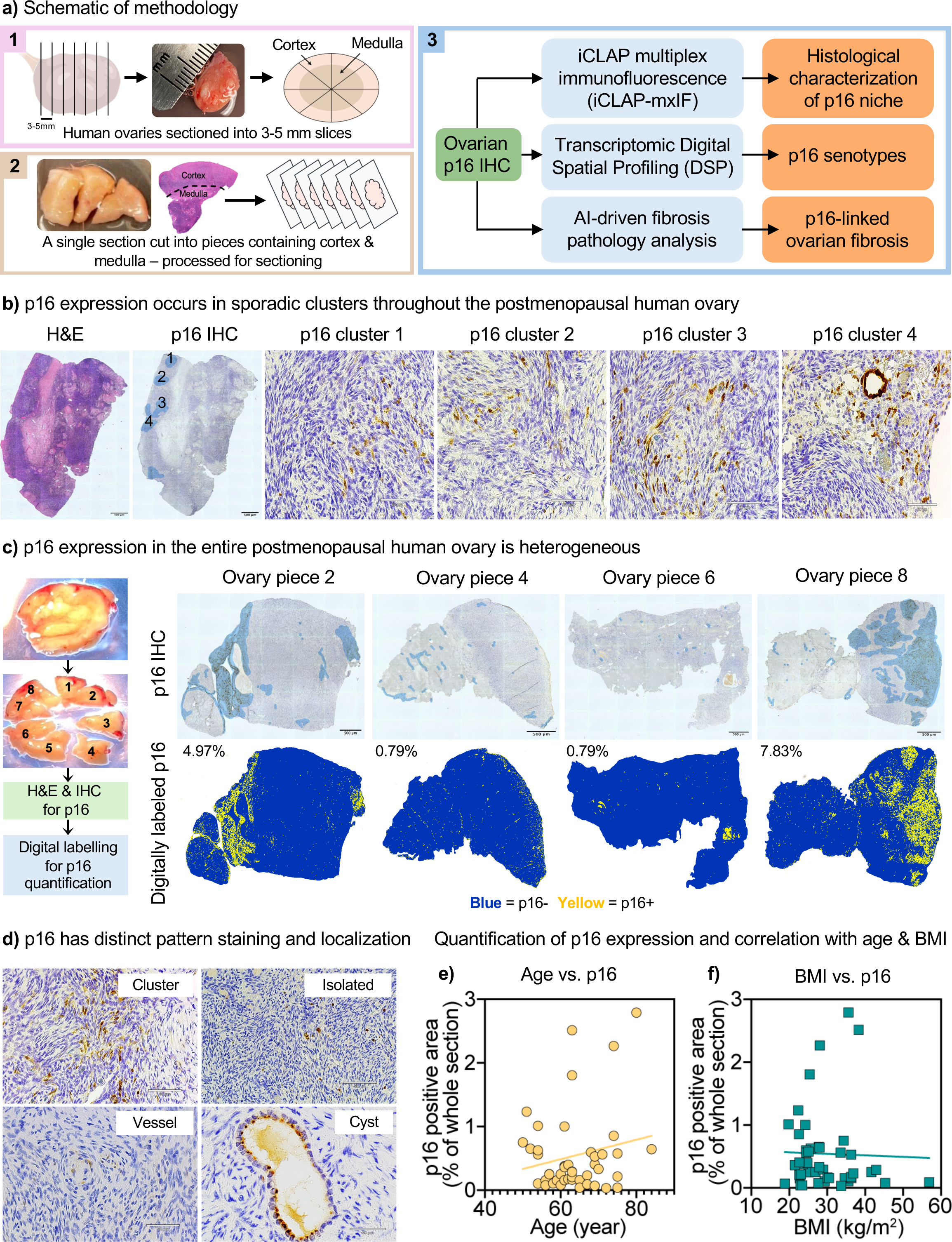
Histological characterization of p16 expression in postmenopausal human ovarian tissue. **a**, Overview of the tissue processing and analysis pipeline. **a1,** Whole postmenopausal ovaries were sliced into 3-5 mm-thick sections. **a2,** Each slice was subdivided into eight pieces containing both cortical and medullary regions. A single tissue piece was formalin-fixed, paraffin-embedded, and sectioned at 5 µm for downstream histological analysis. **a3,** Workflow summary for immunohistochemical staining and image-based quantification of p16^INK4A^ (p16). Blue boxes indicate analytical methodologies; orange boxes indicate corresponding quantitative or visual readouts. **b,** Representative ovarian tissue piece from one participant showing tissue morphology (H&E) and p16 immunostaining with hematoxylin counterstain. Clusters of p16-positive cells are observed sporadically within the tissue. Scale bars: whole section, 500 µm; magnified clusters (1-4), 60 µm. **c,** Heterogeneous distribution of p16 staining in an ovarian section from an 80-year-old participant, subdivided into eight pieces; four pieces (2, 4, 6, and 8) are shown. The top row shows H&E, the middle row shows p16 immunohistochemistry (IHC), and the bottom row shows p16 IHC following digital image labelling. Binary thresholding was applied to identify p16-positive regions (yellow) and p16-negative regions (blue). The percentage of p16-positive area is indicated in the top left of each panel. Scale bar, 500 µm. **d,** Representative p16-IHC images showing clustered or punctate staining and localization to specific structures. Scale bars: cluster and isolated, 60 µm; vessel, 50 µm; cyst, 30 µm. **e-f,** Scatter plots showing the relationship between participant characteristics and tissue-level p16 positivity (n=45). **e,** Percentage of p16-positive staining versus participant age. **f,** Percentage of p16positive staining versus body mass index (BMI) for the same cohort. Simple linear regression lines are shown.

We integrated p16 immunohistochemistry with multiplexed immunofluorescence antibody histology, transcriptomic Digital Spatial Profiling (GeoMx), and AI-guided digital pathology to characterize the p16-positive microenvironment and develop a molecular signature for ovarian p16-positive senescent cells. Whole-transcriptome profiling of 92 spatial regions uncovered a 32-gene p16-associated signature, BuckSenOvary, that distinguished p16-positive regions across cortex and medulla. High-resolution artificial intelligence (AI)-guided analysis of Picrosirius Red-stained tissues revealed increased fibrosis and a shift toward more assembled collagen fibers in p16-positive regions, indicating altered collagen architecture. These regions also showed enrichment for macrophages and myofibroblast-like cells, consistent with an inflammatory-fibrotic feedback loop driving ovarian aging. Together, this study provides the first spatial and molecular characterization of p16-positive senescence-associated fibrosis in the aging native postmenopausal ovary. We identified candidate biomarkers and pathways implicated in ovarian aging, and this study highlights potential molecular targets for senolytic therapies aimed at prolonging or restoring ovarian function. Such strategies may help narrow the healthspan-lifespan gap and preserve systemic health in women.

## Results

### Histological and microenvironment profiling of p16-positive cells in postmenopausal human ovaries

Gynecologic pathologists have long used p16^INK4a^ (p16) protein expression as a surrogate marker in cancer classification and diagnostics for female reproductive tissues, particularly in human papillomavirus (HPV)-driven neoplasia^49–51^. In this context, p16 diagnostic relies on block-positive staining, defined as strong, intense, and continuous nuclear and cytoplasmic staining in basal and parabasal cell layers (Extended Data Fig. 1b). In contrast, patchy weak staining is typically considered background and is non-diagnostic (Fig. 1b and Extended Data Fig. 1a). Interestingly, while this background staining is generally ignored by gynecologic pathologists, it is likely biologically meaningful from an aging and longevity perspective, potentially reflecting tissue stress or cellular senescence, which could act as an anti-cancer failsafe. Indeed, in the aging field, p16 is considered a canonical senescence-associated marker widely used to identify senescent cells^41,52,53^. We set out to characterize these p16-positive cells and their surrounding microenvironment in nonpathological postmenopausal ovarian tissue, where we hypothesized they would be enriched and potentially drive features of ovarian aging.

We first examined the localization and abundance of p16 in ovarian tissue from 45 participants aged 50–84 years. Immunohistochemistry (IHC) revealed that p16 expression within a single ovarian piece of tissue was non-uniform, with sporadic, but distinct clusters of p16-positive cells (Fig. 1b and Extended Data Fig. 2a). These clusters persisted across serial sections (>12 sections, up to 60 μm depth), indicating that they spanned the tissue in three dimensions (Extended Data Fig. 2b). Analysis of a 3-5 mm ovarian cross-section subdivided into eight pieces from a single participant showed that p16 expression was heterogeneous across the ovary, with marked variation in both staining pattern and overall abundance (Fig. 1c and Extended Data Fig. 3a). For example, tissue piece 4 contained 0.79% p16-positive area, whereas tissue piece 8 contained 7.83%. p16 protein expression was observed either as isolated positive cells or as multicellular clusters, predominantly within stromal regions but also in vessels and inclusion cysts (Fig. 1d). IHC staining was digitally labelled using binary thresholding, with p16-positive regions assigned yellow and negative regions blue, enabling quantification of the percentage of p16positive area across tissue sections from all 45 human participants (Extended Data Fig. 3b). When plotted against age, p16 expression trended towards a positive correlation (Fig. 1e), while no correlation was observed with BMI (Fig. 1f), suggesting that age contributed more strongly to increased p16 levels.

To define the microenvironment of p16-positive clusters, we first performed IHC for p21 (CDKN1A; Cyclin-Dependent Kinase Inhibitor 1 A), CD68 (Cluster of Differentiation 68), and α-SMA (alpha-smooth muscle actin) for an initial assessment of expression (Extended Data Fig. 4a). Interestingly, p16-positive cells did not costain with p21, another canonical senescence-associated marker, consistent with emerging studies^31,48^. However, p16-positive regions were enriched for CD68, a macrophage marker, and α-SMA, a marker of myofibroblasts, which are often associated with fibrosis. These observations suggested that p16-positive regions are infiltrated by macrophages and myofibroblasts, which together may elicit alterations in extracellular matrix (ECM) composition and tissue stiffness (Extended Data Fig. 4a).

To characterize these regions more comprehensively, we used iCLAP-mxIF, a multiplex immunofluorescence method optimized for detecting low-abundance proteins (Fig. 2a). This approach enables simultaneous detection of multiple protein markers within a single tissue section. Although ovarian tissue exhibits moderate autofluorescence, particularly from collagen networks and age-related pigments such as lipofuscin, which has historically favored chromogenic IHC and its limitation to one or two markers per section, optimization of iCLAP-mxIF enabled reliable multiplexed staining in the same section. This provided a comprehensive, spatially resolved definition of the p16-positive microenvironment that is not achievable with conventional IHC.

**Figure 2.**
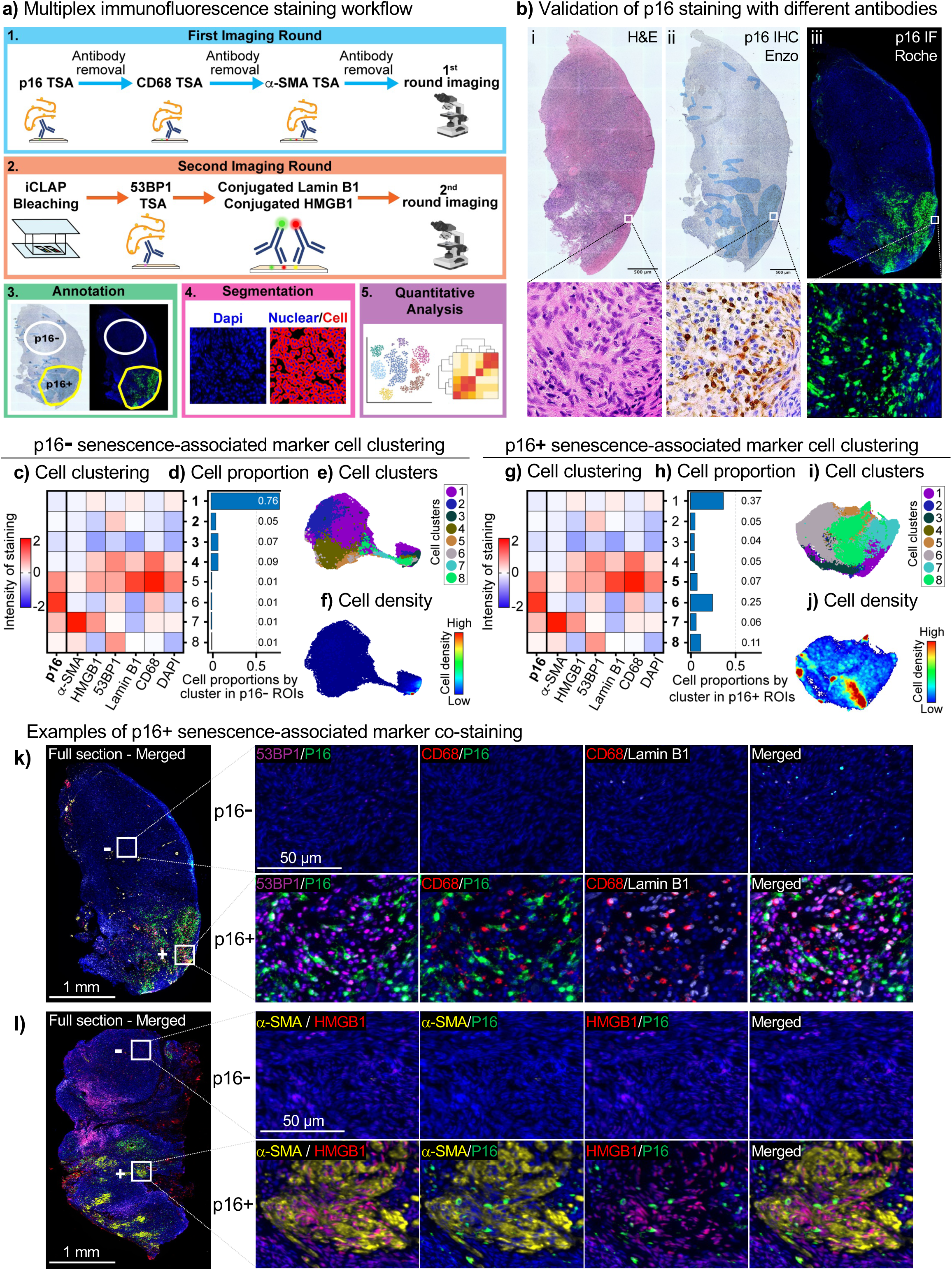
Characterization of p16-positive cells and their niche. **a**, Workflow for multiplex immunofluorescence staining of ovarian tissue. In the first round, p16, CD68, and α-SMA were stained using TSA-based IF. After counterstaining and imaging, slides were bleached and blocked. In the second round, 53BP1 was stained using TSA, followed by direct immunofluorescence detection of Lamin B1 and HMGB1. Images from both rounds were stitched and aligned. Regions were annotated as p16-negative (p16-) or p16-positive (p16+) based on IHC data from Northwestern University. Single-cell segmentation and intensity measurements enabled downstream quantitative analysis. **b,** Validation of p16 immunostaining on human ovary tissue using two distinct antibodies on serial sections. p16 immunohistochemistry (IHC) staining was performed at Northwestern University with a p16 antibody from Enzo Life Sciences Inc. (**bii)**, and immunofluorescence (IF) was performed at Johns Hopkins University with a p16 antibody from Roche Diagnostics on an adjacent slide (**biii**). The adjacent H&E image is shown (**bi**). Representative higher magnification images of H&E and p16 staining are shown below the images of entire sections. **c, g,** Heatmaps of K-means clustering results based on senescence marker intensity profiles from 53865 cells from p16+ regions and 83187 cells from p16-regions. **d, h,** Bar graphs showing the proportion of each cell cluster in p16-(**d**) and p16+ (**h**) regions. **e, I,** UMAPs visualizing the eight cell clusters spatially in p16- (**e**) and p16+ (**i**) regions. **f, j,** UMAPs showing cell density for each cluster in p16- (**f**) and p16+ (**j**) regions. **k,** Example of multiplex IF images from p16- (top row) and p16+ (bottom row) cortical regions. The example highlights cluster 5, p16+/CD68+/Lamin B1+, cluster 6, p16+/53BP1+/CD68+, and cluster 8, p16+/53P1+. Merged channels demonstrate the extent of colocalization between markers. White boxes indicate areas shown at higher magnification in the corresponding panels. **l,** Example of multiplex IF images from p16- (top row) and p16+ (bottom row) cortical regions. The example highlights p16+/a-SMA+/HMGB1+ cluster 7. White boxes indicate regions shown at higher magnification.

The p16 IF staining using an antibody from Roche Diagnostics (Ab 2) was compared and validated against p16 IHC using an antibody from Enzo (Ab 1). Across all four participants, IF consistently recapitulated the same p16-positive regions observed in the IHC sections (Fig. 2b and Extended Data Fig. 4b). This strong concordance between two independent antibodies, detection modalities, and laboratories supports the robustness and specificity of the ovarian p16 signal. After validating p16 staining, we applied the iCLAP-mxIF method (Fig. 2a). We used six senescence-associated markers: p16, α-SMA, HMGB1 (High Mobility Group Box 1, 53BP1 (p53-binding protein 1), Lamin B1, and CD68. Regions of p16-negative and p16-positive expression were annotated across tissue sections, and single-cell segmentation was performed to generate marker intensity profiles (Fig. 2a). While segmentation can introduce occasional bleed-through between neighbouring cells, marker intensity profiles were obtained from 83,187 cells in p16-negative regions and 53,865 cells in p16-positive regions.

We next applied minimal clustering to the single-cell marker profiles, which resolved eight populations based on expression of the six senescence-associated markers (Fig. 2c, 2g, and Supplemental Table 2). In the heatmaps, each row represents a cluster, and each column a marker, with color indicating relative staining intensity. In p16-negative regions, most cells belonged to clusters 1-4, with cluster 1 alone comprising 76% of cells, and clusters 2-4 each contributing 5-9% (Fig. 2c-e). The p16-negative regions were relatively homogeneous in composition, as reflected in the cell density UMAP (Fig. 2f). Clusters 1-3 exhibited uniformly low expression of all six markers, whereas Cluster 4 displayed low levels of p16 and α-SMA but relatively high expression of HMGB1, 53BP1, Lamin B1, and CD68 (Fig. 2c). This pattern may represent activated or stressed macrophages, or cells with features of DNA damage and stress consistent with a pre-senescent state in proximity to infiltrating macrophages.

In p16-positive regions, the same eight clusters were present, but their relative abundance shifted markedly (Fig. 2g-i). Clusters 5-8 (p16-high populations) collectively accounted for 49% of all cells, with Cluster 6 being the most abundant (25% of cells), whereas p16-low clusters 1-4 together represented 51% and were still dominated by cluster 1 (37%) (Fig. 2g-i). Unlike the relative homogeneity of p16negative regions, p16-positive regions displayed marked cellular heterogeneity, as shown by the cell density UMAP (Fig. 2j). Cluster 5 expressed all six markers and accounted for 7% of cells, but showed minimal CD68 colocalization, suggesting segmentation bleed-through from neighbouring macrophages (Fig. 2k and Extended Data Fig. 4c). Importantly, this analysis confirmed that p16-positive cells in ovarian tissue are not exclusively macrophages but include a substantial population of CD68 negative stromal cells. Cluster 6, the most abundant p16-positive cluster, coexpressed p16, 53BP1, and CD68, suggesting a senescent-like macrophage subset or a non-macrophage pre-senescent/stromal population (Fig. 2g-h and Extended Data Fig. 4c). Cluster 8, positive for p16, α-SMA, and 53BP1, showed a myofibroblast-like senescent phenotype (Fig. 2l), a population of particular interest, as it may drive the fibrotic remodelling in the aging ovary. Further studies will be required to delineate these candidate senescent populations more precisely. To move beyond protein-level marker profiling and define the broader molecular programs operating in these p16-positive niches, we next applied spatial transcriptomics.

### Spatial transcriptomic characterization and signature derivation of p16positive senescent cells

To define the molecular signature of p16-positive compared to p16-negative regions, we profiled these areas using spatial transcriptomics. An ovarian tissue piece from an 80-year-old participant was serially sectioned at a thickness of 5 µm each, with alternating sections designated for either p16 IHC or GeoMx Digital Spatial Profiling (DSP) (Fig. 3a). p16 IHC-stained slides were annotated for p16-positive clusters and used to define p16-positive and p16-negative regions of interest (ROIs) on adjacent DSP sections (Fig. 3a-b and Extended Data Fig. 5b). This approach was feasible because p16-positive clusters spanned three dimensions within the ovarian tissue (Extended Data Fig. 2b, 5a). Following acquisition and sequencing of GeoMx slides, transcriptomic analysis was performed across three independent tissue sections, enabling the identification of differentially expressed genes and the derivation of p16associated transcriptomic signatures (Fig. 3c and Extended Data Fig. 6).

**Figure 3.**
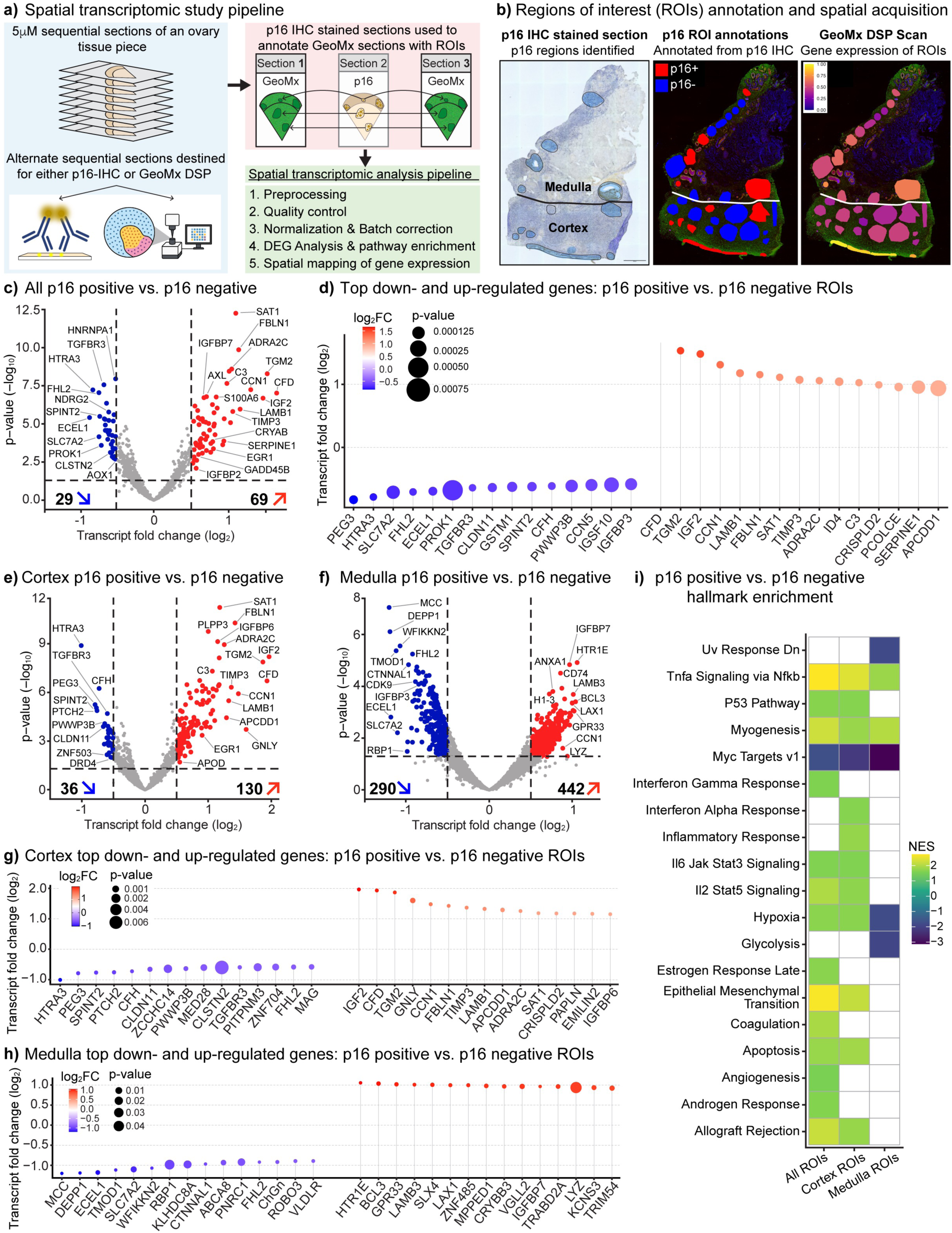
Spatial transcriptomic profiling of p16-positive and p16-negative regions in postmenopausal ovary. **a**, Overview of the spatial transcriptomic study design. Serial 5 mm paraffin sections from a single ovarian tissue piece of an 80year-old participant were allocated for GeoMx Digital Spatial Profiling (DSP), p16 immunohistochemistry (IHC), or left unstained for future analysis. p16 IHC-stained sections were used to annotate adjacent sections for DSP, enabling selection of p16positive (p16+) and p16-negative (p16-) regions of interest (ROIs). Sections 1, 3, and 7 were processed for transcriptomic analysis. **b,** Example of ROI identification and transfer. Section 2, stained for p16 by IHC, was used to annotate p16+ and p16-ROIs across the ovarian tissue section, including cortical, medullary, vascular, and cystic regions. These annotations were transferred to the adjacent GeoMx DSP section (section 1) for targeted transcriptomic profiling. **c,** Differential gene expression analysis of all p16+ versus p16-ROIs across sections 1, 3, and 7 (n = 92). Volcano plot shows significantly up-and downregulated genes (adjusted *P* < 0.05; log_2_ fold change > 0.5). **d,** Dot plot showing the top 15 most upregulated and downregulated transcripts in p16+ versus p16-ROIs across all regions. **e,** Subanalysis of cortical ROIs (n=71). Volcano plot shows differentially expressed genes between p16+ and p16-cortex-only regions (adjusted *P* < 0.05; log_2_ fold change > 0.5). **f,** Sub-analysis of medullary ROIs (n=21). Volcano plot shows differentially expressed genes between p16+ and p16-medulla-only regions (adjusted *P* < 0.05; log_2_ fold change > 0.5). **g,** Dot plot showing the top 15 most upregulated and downregulated transcripts in cortex p16+ versus p16-ROIs. **h,** Dot plot showing the top 15 most upregulated and downregulated transcripts in medulla p16+ versus p16-ROIs. **i,** Pathway enrichment analysis comparing p16+ versus p16-ROIs across all tissue (All), cortex only, and medulla only groups. Normalized enrichment scores (NES) are shown for selected gene sets.

We performed three comparisons: i) all p16-positive versus all p16-negative regions of interest (ROIs) within the ovary, and subsequently investigating the functionally distinct ovarian tissue compartments, cortex and medulla, comparing ii) cortex p16positive versus cortex p16-negative ROIs, and iii) medulla p16-positive versus medulla p16-negative ROIs. Across all ROIs, 29 genes were downregulated, and 69 were upregulated (adjusted P <0.05, log_2_ fold change >0.5) (Fig. 3c and Supplementary Table 3). The top 15 downregulated genes included IGFBP3, CCN5, CFH, SPINT2, and TGFBR3, regulators of growth-factor, complement, and TGF-β signaling that have been implicated in restraining excessive proliferation and inflammation (Fig. 3d). The top 15 upregulated genes comprised canonical SASP factors (SERPINE1, CFD, C3, and CCN1) as well as extracellular matrix (ECM)remodeling genes (TGM2, LAMB1, FBLN1, and PCOLCE) (Fig. 3d and Extended Data Fig. 7a-b).

In ovarian cortical ROIs, 36 genes were downregulated, and 130 were upregulated, with the top 15 down-and up-regulated genes largely overlapping with the whole-tissue comparison (Fig. 3e, 3g, and Supplementary Table 4). This suggested that the ovarian cortex carried the core senescence/fibrosis signature (Fig. 3i). By contrast, ovarian medullary ROIs showed a far broader transcriptional response, with 290 downregulated and 442 upregulated genes (Fig. 3f and Supplementary Table 5). The top 15 down-and up-regulated genes in the medulla did not overlap with those from the whole tissue or cortex analyses and instead reflected pro-inflammatory, immune surveillance, and DNA damage-response pathways (Fig. 3h).

Pathway enrichment analysis revealed broad activation of hallmark senescence programs in p16-positive ROIs, including TNFα signaling via NF-κB, interferon α/γ response, IL6-JAK-STAT3, and IL2-STAT5 signaling, as well as p53-mediated growth arrest, along with suppression of proliferative Myc targets (Fig. 3i). These findings indicate activation of SASP-like inflammatory signaling and immune crosstalk within p16-positive regions. Simultaneously, we observed enrichment of epithelialmesenchymal transition (EMT), coagulation, angiogenesis, and hypoxia pathways, suggesting that senescent cells in the aging ovary promote fibrotic remodeling and vascular adaptation, features that are consistent with a pro-fibrotic senescence phenotype (Fig. 3i). The cortical p16-positive regions largely recapitulated these senescence-associated pathways, showing strong activation of NF-κB, interferon, and EMT programs (Fig. 3i). In contrast, medullary p16-positive regions showed selective enrichment of glycolysis and hypoxia pathways, with suppression of Myc target and UV response programs, suggesting a metabolically repressed, stress-adapted senescence state (Fig. 3i). These compartment-specific signatures suggest that ovarian senescence has distinct transcriptional states shaped by local microenvironmental cues, ranging from inflammatory-fibrotic in the cortex to hypoxia-adaptive in the medulla.

After examining overall transcriptomic differences between p16-negative and p16positive regions, we next sought to derive gene signatures that could robustly define p16-positive regions and potentially enable their use for unsupervised mapping. Because each analytical approach captures distinct aspects of the p16-associated transcriptomic landscape and can yield partially non-overlapping gene sets, we applied multiple complementary methods to derive convergent signatures that robustly define p16-positive regions across cortical and medullary compartments. We first identified significantly differentially expressed genes (DEGs; P <0.05) between p16-positive and p16-negative regions across all ROIs, cortical ROIs, and medullary ROIs, considering both upregulated and downregulated DEGs. The overlap among these three comparisons was then assessed to identify shared signatures independent of ovarian region (Fig. 4a, c). This analysis yielded 90 shared downregulated DEGs, which we designated *Signature 1* (Fig. 4a-b and Extended Data Fig. 8a). These genes reflected broad suppression of cell-cycle drivers (CCNB1IP1, CCN5, TGFBR3), metabolic regulators (SLC7A2, RBP1, PDK4) (Fig. 4b), and, most prominently, ribosomal and translational factors (RPL/RPS family members, EIF4B) (Extended Data Fig. 8a). Pathway enrichment confirmed marked depletion of translation, ribosome biogenesis, and rRNA processing, consistent with the established biology of senescent cells, which downregulate proliferation and protein synthesis while maintaining SASP activity (Extended Data Fig. 8b)^54–56^.

**Figure 4.**
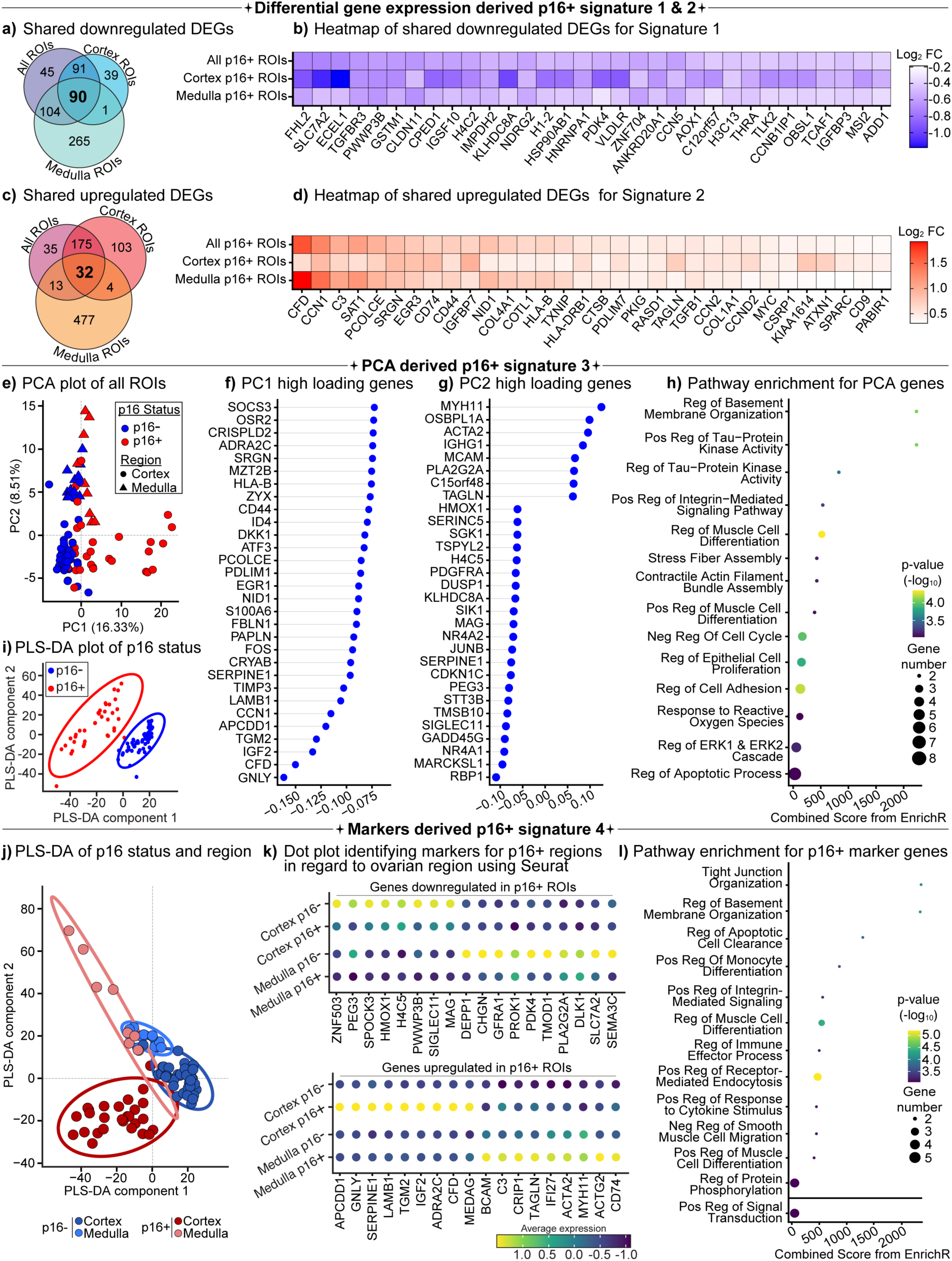
Derivation of p16-associated transcriptomic signatures to map senescent cells. Multiple analytical approaches were applied to identify robust p16associated gene signatures for subsequent evaluation of p16-senescence-associated specificity and precision. **a,** Venn diagram showing overlap of downregulated differentially expressed genes (DEGs; *P* < 0.05) between p16positive (p16+) and p16-negative (p16-) regions of interest (ROIs) across three comparisons: all ROIs combined (All), cortex-only ROIs, and medulla-only ROIs. **b,** Heatmap of 32 shared downregulated DEGs common to all three comparisons, defined as *Signature 1*. **c,** Venn diagram showing overlap of upregulated DEGs (*P* < 0.05) across the same three comparisons. **d,** Heatmap of 32 shared upregulated DEGs common to all three comparisons, defined as *Signature 2*. **e,** Principal component analysis (PCA) of all ROIs, colored by p16 status (blue, p16-; red, p16+) and annotated by anatomical region (cortex or medulla). **f,** Dot plot of top-loading genes from principal component 1 (PC1), which separates ROIs by p16 status, defining *Signature 3*. **g,** Dot plot of top-loading genes from principal component 2 (PC2), which separates ROIs by anatomical region. **h,** Pathway enrichment analysis of genes contributing to PC1, highlighting biological processes associated with p16 status. **i,** Partial least squares discriminant analysis (PLS-DA) of ROIs based on p16 status alone (p16+ vs p16-). **j,** PLS-DA incorporating both p16 status and anatomical region (cortex vs medulla). **k,** Dot plot of average gene expression for marker genes distinguishing p16 status and region, identified using the Seurat FindAllMarkers() function. Upregulated markers defining p16+ status constitute *Signature 4*. **l,** Pathway enrichment analysis of upregulated marker genes from *Signature 4*. *Signatures 1-4* are later validated in Fig. 5.

We next identified 32 upregulated DEGs shared across all, cortical, and medullary comparisons, which we designated *Signature 2* (Fig. 4c). These genes reflected activation of inflammatory and fibrotic pathways, including complement factors (CFD, C3), extracellular matrix components (CCN1, PCOLCE, COL1A1/2), and regulators of SASP (TGFB1, TXNIP) (Fig. 4d). Pathway enrichment analysis demonstrated positive regulation of ERK/MAPK signaling, vascular growth factor production, extracellular matrix organization, and monocyte differentiation, alongside suppression of apoptosis. Collectively, these data indicated that p16-positive regions adopted a pro-inflammatory, pro-fibrotic, and survival-oriented state consistent with senescence-associated secretory activity and tissue remodeling (Extended Data Fig. 8c).

As a second approach, we applied principal component analysis (PCA) to reduce transcriptomic complexity and identify the largest sources of variation underlying p16-associated profiles. PCA distinguished p16-positive from p16-negative regions along PC1, which accounted for 16.3% of the variance and captured the senescence transcriptional profile, designated *Signature 3* (Fig. 4e). PC1 loadings were enriched for SASP and ECM regulators (SERPINE1, TIMP3, CCN1, LAMB1, FBLN1, TGM2, IGF2, CFD) and immune mediators (CD44, HLA-B, SRGN). Pathway analysis confirmed downregulation of cell-cycle progression and upregulation of extracellular matrix organization, integrin-mediated adhesion, oxidative stress response, and ERK/MAPK signaling (Fig. 4f, h). In contrast, PC2 (8.5% variance) regionally separated cortical from medullary ROIs and was driven by smooth muscle and stress-response genes (MYH11, ACTA2, TAGLN, HMOX1), highlighting regional heterogeneity in senescent signatures (Fig. 4e, g). A supervised partial least squares-discriminant analysis (PLS-DA) also confirmed clear separation of p16positive from p16-negative regions, driven by ECM regulators, immune mediators, and stress-response genes (TIMP3, CCN1, LAMB1, C3, TXNIP) (Fig. 4i and Extended Data Fig. 8d, e). Pathway enrichment showed downregulation of proliferative programs (Myc, KRAS, UV response) and upregulation of inflammatory, apoptotic, and EMT pathways, consistent with a senescence-associated remodeling signature (Extended Data Fig. 8f-g).

For the third approach to define a p16 signature, we found that when performing a partial least squares discriminant analysis (PLS-DA), a supervised approach using both p16 status and ovarian region, it demonstrated that p16 status and ovarian region could be clearly separated by transcriptomic profiles (Fig. 4j). To derive a regionally informed p16 signature, we applied Seurat to identify marker genes distinguishing p16-positive regions across cortex and medulla (Fig. 4k). *Signature 4* was defined by upregulated genes including ECM regulators (SERPINE1, LAMB1), immune mediators (C3, CD74), and stress-response factors (ACTA2) (Fig. 4k). Pathway enrichment confirmed activation of extracellular matrix organization, integrin signaling, immune effector processes, and apoptotic regulation, consistent with a senescence-associated remodeling program (Fig. 4l).

### Evaluation of transcriptomic signatures to identify and map p16-positive regions

We identified p16-associated gene signatures (Signatures 1-4; each comprising 30-90 genes) condensed from ∼3000 differentially expressed genes (Supplementary Table 6). We then evaluated the accuracy with which each signature distinguished p16-positive from p16-negative regions by applying them in an unsupervised manner to 92 ROIs from three ovarian sections. For each signature, we calculated a UCell enrichment score, referred to as the “p16 mapping score”, and overlaid these scores onto tissue sections to visually assess mapping performance. Signature 1, based on shared downregulated DEGs (Fig. 4a-b and Extended Data Fig. 8a), significantly distinguished p16-positive from p16-negative regions (*p=1.4×10^-^*^9^) (Fig. 5a-b and Extended Data Fig. 9a). Signature 2, derived from shared upregulated DEGs (Fig. 4c-d), also significantly identified p16-positive regions (*p=1.1×10^-^*^9^) (Fig. 5d-e and Extended Data Fig. 9a). Similarly, Signature 3, generated from PCA analysis (Fig. 4e-f), and Signature 4, generated from Seurat marker analysis (Fig. 4j-k), both significantly identified p16-positive regions (*p=1.5×10^-^*^5^ *and p=4×10^-^*^8^) (Fig. 5g-h, j-k, and Extended Data Fig. 9a).

**Figure 5.**
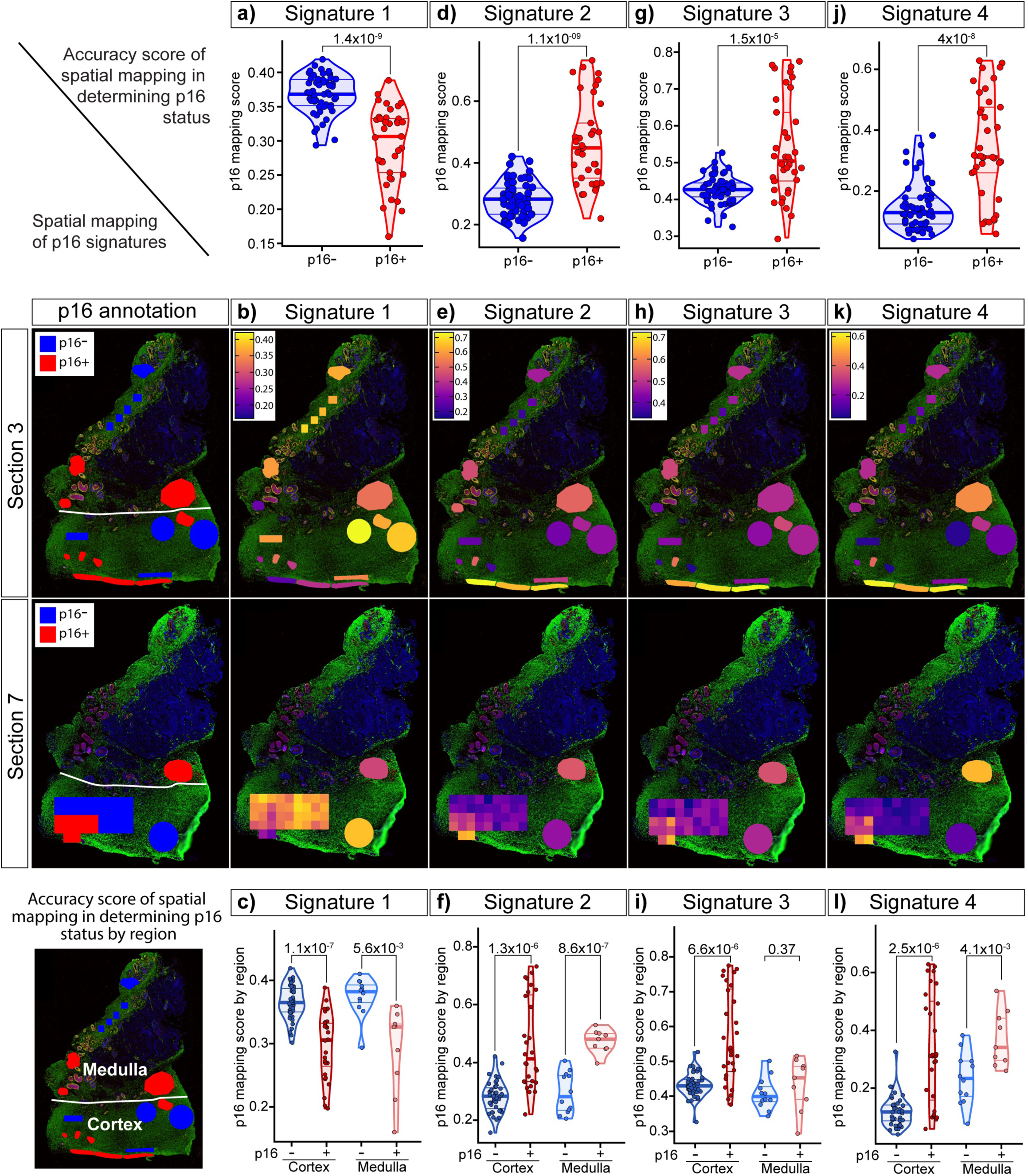
Evaluation of p16-associated transcriptomic signatures for spatial mapping of p16-positive regions. The four p16-associated gene signatures (Signatures 1-4), derived from 92 regions of interest (ROIs) across three ovarian sections, were assessed for their ability to distinguish p16-positive (p16+) from p16negative (p16-) regions and to spatially map senescent cells. **a, d, g, j,** Violin plots showing UCell-derived enrichment scores (“p16 mapping scores”) for each signature in ROIs annotated as p16-(blue) or p16+ (red). **b, e, h, k,** Unsupervised spatial mapping of each signature to ovarian sections 3 and 7 using the SpatialOmicsOverlay package. Enrichment scores are overlaid onto tissue architecture; far-left panels indicate reference p16 status annotation for each ROI (p16-, blue; p16+, red). **c, f, i, l,** Violin plots showing UCell enrichment scores for each signature stratified by anatomical region (cortex vs medulla) and p16 status. This analysis tests whether signatures discriminate p16+ regions independently of tissue region. Violin plots were analyzed using two-sided t-tests; *P* < 0.05 was considered significant.

When stratified by ovarian region, differences in signature performance emerged. Signature 3 was able to significantly identify p16-positive regions in the cortex (*p=6.6×10^-^*^6^) but failed to do so in the medulla (*p=0.37*) (Fig. 5i). By contrast, Signatures 1, 2, and 4 significantly identified p16-positive regions in both cortex and medulla (Fig. 5c, f, i). Signature 2 showed the most significant performance in distinguishing p16 status amongst all ROIs (*p=1.1×10^-^*^9^), ROIs in cortical regions (*p=1.3×10^-^*^6^), and ROIs in medullary regions (*p=8.6×10^-^*^7^) (Fig. 5d, e, f). This leverages Signature 2 as the strongest transcriptomic signature that defines a p16positive region regardless of ovarian regional location.

We next evaluated external senescence gene sets against our ovarian signatures. The curated “SenMayo” gene set^57^ (Supplementary Table 6), which has been widely used to detect senescent cells, significantly distinguished p16-positive from p16negative regions (Extended Data Fig. 9a-b) and performed consistently across ovarian regions (Extended Data Fig. 9c). However, its performance was weaker to our internally derived Signature 2 (Fig. 5f). We also tested a proteomic SASP signature that we had previously generated by inducing senescence in ovarian tissue with doxorubicin^58^ (Supplementary Table 6). This ovary-specific SASP signature significantly identified p16-positive regions (Extended Data Fig. 9d) but showed weaker overall performance and failed to discern p16 status when stratified by ovarian region (Extended Data Fig. 9e), consistent with it being a secreted proteomic signature. Together, these analyses support the conclusion that p16-positive regions represent bona fide senescent regions within the aging ovary.

Having established signatures capable of distinguishing p16-positive regions, we next evaluated their ability to map these regions in native ovarian tissue. Tissue sections from the ovarian tissue of two participants, aged 67 (participant 1) and 71 (participant 2) years old, were analyzed. Each section was gridded into 400 × 400 μm boxes and profiled using GeoMx DSP. Signatures 2 and 4 were then applied to assess whether they could identify p16-positive regions, with validation performed by cross-referencing to p16 IHC staining on adjacent sections. In participant 1, hotspots defined by Signatures 2 and 4 (Extended Data Fig. 10a) overlapped with p16-positive clusters 1 and 2 detected by IHC (Extended Data Fig. 10b). Similarly, in participant 2, hotspots detected by Signatures 2 and 4 (Extended Data Fig. 10c) coincided with p16-positive clusters 1-3 identified by IHC (Extended Data Fig. 10d). These findings provided proof-of-principle that our transcriptomic signatures can spatially map p16-positive regions in an unsupervised manner. However, the resolution of GeoMx DSP limited precise cellular mapping, and future evaluation of these signatures will require single-cell spatial technologies such as CosMx. Given that these p16-associated signatures and pathway analyses consistently highlighted extracellular matrix remodelling and fibrosis, we next sought orthogonal histological evidence for fibrotic changes in p16-positive regions.

### AI-driven pathology reveals fibrosis enrichment in p16-positive ovarian regions

Spatial transcriptomic analyses revealed enrichment of extracellular matrix (ECM) remodeling and fibrosis-associated pathways in p16-positive regions (Fig. 3, 4, and Extended Data Fig. 7). To directly assess transcriptional matrisome changes, we examined gene expression using the curated “Matrisome Project” database^59^. Several collagens, ECM glycoproteins, proteoglycans, regulators, and secreted factors were enriched in p16-positive ROIs (Fig. 6a), suggesting that p16-positive cells may shape the ECM microenvironment to promote fibrotic remodelling. Furthermore, we noticed that p16-positive regions coincided with sclerotic areas when evaluating histological sections (Fig. 6b). To validate this at the tissue level, we performed picrosirius red staining in ovarian sections from ten participants, followed by high-resolution slide scanning and AI-based image analysis (FibroNest) (Fig. 6c). This approach quantified over 300 quantitative fibrosis traits (qFTs), which were aggregated into 32 phenotypic traits across matched p16-negative and p16-positive ROIs, encompassing bulk collagen deposition, fiber morphology, and architectural complexity (Fig. 6d-e and Supplementary Table 7).

**Figure 6.**
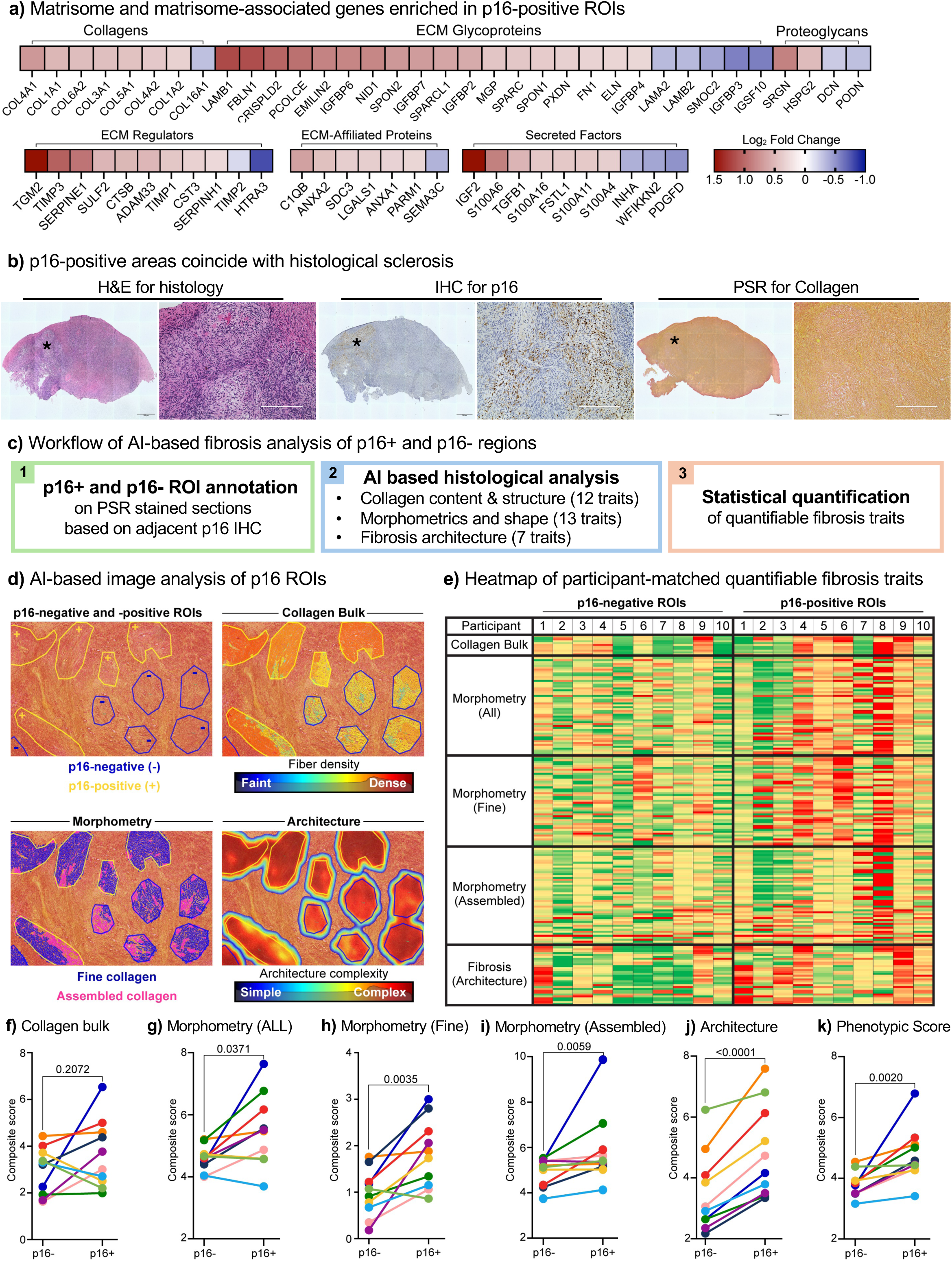
AI-driven digital pathology confirms fibrotic enrichment in p16positive ovarian regions. Fibrosis-associated features were quantified in participant-matched p16-positive (p16+) and p16-negative (p16-) tissue regions from ten participants to assess the impact of p16 expression within the ovarian stromal niche, integrating transcriptomic matrisome signatures with high-resolution collagen imaging. **a,** Heatmap of matrisome and matrisome-associated genes significantly enriched in p16+ regions of interest (ROIs) relative to p16-ROIs, represented as log_2_ fold change. **b,** Images of Hematoxylin and eosin (H&E), p16-IHC and eosin, and picosirius red staining that highlight co-localization of histological sclerosis with p16positive regions. **c,** Workflow summarizing the AI-based fibrotic analysis pipeline using the FibroNest platform (PharmaNest) applied to p16-annotated sections. **d,** Representative picosirius red–stained ovarian sections from ten participants, annotated for p16 status and analyzed for quantitative collagen traits including bulk, morphometry, and architecture. **e,** Heatmap of fibrosis severity for matched p16+ and p16-ROIs across participants. Severity is color-coded from green (fine collagen, minimal fibrosis) to red (dense, complex collagen, maximal fibrosis). Over 300 quantitative fibrosis traits (qFTs) were measured and aggregated into three composite categories: collagen bulk (12 traits), morphometry (13 traits), and architecture (7 traits). **f–k,** Quantification of normalized composite fibrosis scores derived from **e**. Scores range from 0 to 10, with higher values indicating greater fibrosis. Data distribution was assessed using the Shapiro–Wilk normality test: panels **f**, **h**, and **j** passed and were analyzed using paired t-tests; panels **g**, **i**, and **k** were analyzed using Wilcoxon matched-pairs signed rank tests. *P* < 0.05 was considered significant.

Heatmap visualization revealed that p16-negative ROIs generally scored lower (green, less fibrosis), whereas p16-positive ROIs scored higher (red, more fibrosis) (Fig. 6e and Supplementary Table 7)). Composite trait scoring demonstrated significant increases in fine collagen (*p=0.0035*), assembled collagen (*p=0.0059*), and fibrosis architecture (*p<0.0001*), yielding a higher overall phenotypic fibrosis score in p16-positive regions (*p=0.0020*) (Fig. 6g-k). Bulk collagen content, however, did not differ significantly (*p=0.21*) (Fig. 6f), consistent with our independent whole tissue (irrespective of p16 status) picrosirius red quantification (Extended Data Fig. 11). These findings indicated that p16-positive regions are fibrotically remodelled not by an increase in total collagen, which is already elevated in postmenopausal ovaries, but by alterations in ECM organization, such as fiber architecture, crosslinking, and stromal stiffness, features more sensitively captured by AI-based fibrosis scoring than by bulk collagen quantification.

### The p16 senotype BuckSenOvary reflects senescence, inflammation, and fibrosis in the aging ovary

Taken together, our multiplex protein, spatial transcriptomic, and AI-based pathology analyses indicate that p16-positive regions in the postmenopausal human ovary constitute a senescent stromal niche characterized by inflammation and fibrosis. While p16 is widely recognized as a canonical marker of cellular senescence, our analyses show that p16-positive regions in the human ovary are enriched for senescence-associated pathways (Fig. 3i), and that the curated SenMayo senescence gene set^57^ robustly identifies these regions (Extended Data Fig. 9a,b). In parallel, we recently developed a postmenopausal human ovarian explant culture model in which low-dose doxorubicin induces cellular senescence while preserving tissue viability, and multi-omics profiling (snRNA-seq and proteomics) defined an ovary-specific senescence signature (“ovarian senotype”) comprising 26 overlapping transcriptomic and proteomic targets, several of which were validated in native aged ovarian tissue^31^. To determine whether p16-positive regions reflect similar transcriptionally senescent cell states, we compared their transcriptomes with those of doxorubicin-treated postmenopausal ovarian explants from our previous study^31^. We observed strong concordance across both cortical and medullary compartments, with many senescence-associated genes changing in the same direction in the two datasets (Fig. 7a). Key overlapping transcripts included CCN1, SAT1, CD44, NID1, IGFBP7, and PCOLCE, all upregulated in p16-positive regions and in doxorubicin-treated explants (Fig. 7a). These genes regulate fibroblast activation, stress adaptation, and secretory function, features consistent with the senescence-associated secretory phenotype (SASP) and matrix remodeling. In contrast, structural and metabolic genes such as VIM and NRK were decreased in both models, indicating a shift toward a secretory-fibrotic stromal state. Interestingly, seven genes, representing 22 % of the “p16 Signature 2” (BuckSenOvary), overlapped with the doxorubicin-induced senescence signature (Fig. 7a). Together, these findings link experimentally induced and endogenous senescence programs, supporting that p16-expressing cells in the aging ovary are transcriptionally senescent stromal populations.

**Figure 7.**
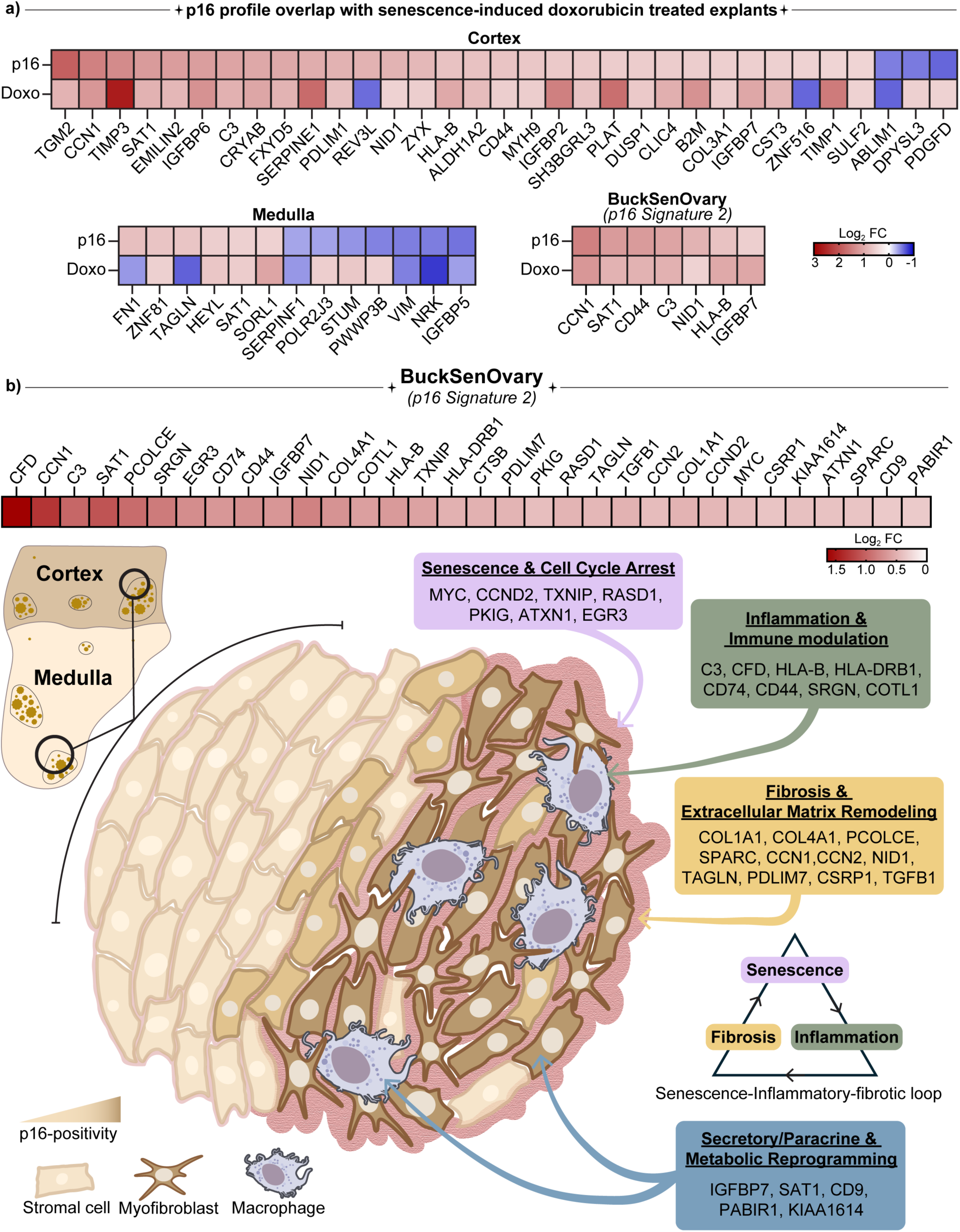
p16 senotype BuckSenOvary reflects senescence, inflammation, and fibrosis in the aging ovary. Transcriptomic comparison of doxorubicin-induced senescence in human ovarian explants and p16-positive regions in native human ovary reveals overlapping gene expression profiles across cortical and medullary compartments, including the p16-associated signature, BuckSenOvary. Heatmaps show log_2_ fold changes of overlapping genes (*P* < 0.05, log_2_ fold change > 0.5) identified in both datasets. The BuckSenOvary senotype comprises four coordinated biological modules: **senescence and cell-cycle arrest** (MYC, CCND2, TXNIP, RASD1, PKIG, ATXN1, EGR3), **inflammation and immune modulation** (C3, CFD, HLA-B, HLA-DRB1, CD74, CD44, SRGN, COTL1), **fibrosis and extracellular matrix** remodeling (COL1A1, COL4A1, PCOLCE, SPARC, CCN1, CCN2, NID1, TAGLN, PDLIM7, CSRP1, TGFB1), and **secretory/paracrine and metabolic reprogramming** (IGFBP7, SAT1, CD9, PABIR1, KIAA1614). The schematic depicts the transition from histologically homogeneous p16-negative regions to heterogeneous p16-positive zones enriched for senescent stromal cells, macrophage infiltration, and complex fibrotic architecture. Together, these findings define a self-reinforcing senescence-inflammation-fibrosis feedback loop represented by the p16-linked senotype BuckSenOvary, which underlies ageassociated ovarian remodeling.

Among all the transcriptomic p16 signatures evaluated, Signature 2 most significantly distinguished p16-positive regions across all ROIs (*p=1.1×10^-^*^9^), and regionally within both the cortex (*p=1.3×10^-^*^6^), and the medulla (*p=8.6×10^-^*^7^) (Fig. 5d-f). This consistency across compartments establishes Signature 2 as a defining molecular fingerprint of p16-positive senescence in the human ovary, which we have termed “BuckSenOvary” (Fig. 7b). The BuckSenOvary senotype comprises four interconnected biological modules that together define the transcriptional landscape of p16-positive ovarian regions (Fig. 7b). The first is a senescence and cell-cycle regulatory module (MYC, CCND2, TXNIP, RASD1, PKIG, EGR3, and ATXN1) that reflects altered proliferative signaling, oxidative stress responses, and Ras-cAMP pathways characteristic of p16-mediated growth arrest, and pointing to reinforced stress surveillance through TXNIP-dependent p53 activation and emerging links between ATXN1 and DNA damage–associated stress granules. The inflammatory and immune response module, enriched for SASP-associated mediators (C3, CFD, HLA-B, HLA-DRB1, CD74, CD44, SRGN, COTL1), captures complement activation, antigen presentation, and macrophage-stromal signaling that connect senescence to chronic inflammation and tissue remodeling. The fibrosis and extracellular matrix remodeling module contains matrisome components (COL1A1, COL4A1, PCOLCE, SPARC, CCN1, CCN2, NID1, TAGLN, PDLIM7, CSRP1, TGFB1) that promote collagen deposition, cytoskeletal remodeling, and matrix organization, features typical of fibroblast-like senescent cells. Finally, the secretory and metabolic adaptation module (IGFBP7, SAT1, CD9, PABIR1, and KIAA1614) reflects oxidative stress adaptation, early-response transcriptional control, and vesicle-mediated communication. Among these, IGFBP7 stands out as a key secretory factor, consistent with its role in fibroblast senescence and the SASP. Together, these modules define BuckSenOvary as a coordinated stress-response program that integrates cell-cycle arrest, inflammation, and fibrotic remodeling, engaging in a senescence-inflammatory-fibrotic loop in the aging ovary (Fig. 7b).

## Discussion

Cellular senescence, a hallmark of aging across multiple tissues, has remained poorly defined in the human ovary. This gap stems from the challenges of identifying senescent cells *in vivo*, the lack of senescence biomarkers, and the scarcity of healthy, non-pathological ovarian tissue. We previously optimized a postmenopausal ovarian explant culture model, inducing senescence with low-dose doxorubicin, enabling a multiomic definition of an ovarian senotype^31^. Building on this, we conducted integrative analyses using the canonical senescence-associated marker p16^INK4a^ in native postmenopausal ovarian tissue. Together, these analyses revealed discrete senescent niches: p16-positive clusters were rare, spatially heterogeneous, and enriched within stromal, vascular, and cystic regions, often in proximity to macrophages. Multiplex protein imaging further resolved these niches into distinct senescent-like macrophage and myofibroblast populations alongside stressed, p16low stromal cells, highlighting cellular heterogeneity within the p16-positive microenvironment. Spatial transcriptomic profiling showed that p16-positive regions were characterised by suppression of cell-cycle and translational machinery, alongside activation of inflammatory, immune, and extracellular matrix (ECM) remodelling programs. Among four tested transcriptomic signatures, our novel ovary-specific BuckSenOvary signature (*Signature 2,* 32-gene ovary-derived) most significantly distinguished p16-positive from p16-negative regions compared with the 125-gene bone-derived SenMayo senescence gene set, highlighting the need for elucidating organ-specific senescence signatures (senotypes).Notably, these signatures could also map p16-positive regions in an unsupervised manner in independent tissues. AI-driven pathology further demonstrated that p16-positive regions were not marked by increased bulk collagen, but rather by qualitative changes in ECM architecture and organisation. Collectively, these findings position p16-positive microenvironments as fibro-inflammatory, potentially contributing to ovarian ageing and downstream systemic health.

p16^INK4a^ has become the most widely used marker for identifying senescent cells in both research and clinical settings^41,60^. Its advantages include its established link to cell-cycle arrest through CDK4/6 inhibition^44,61–63^, its increased expression with age across various tissues^39,47,64^, and its reliability in histological assays where other senescence markers are less consistent^65^. In the ovary, mouse studies have shown that p16 expression increases with age, especially in stromal regions, which matches our observations in human tissue^39,66^. Transgenic reporter models such as p16^Ink4a^luciferase and p16^Ink4a–^GFP further confirm that p16-positive cells accumulate with age and reproductive aging. These cells contribute to follicular decline and stromal fibrosis^66–70^. However, systematic studies in humans are limited due to the scarce availability of healthy ovarian tissue. Transcriptomic analyses generally have not detected age-related increases in p16 expression in the human ovary, though elevated p21 transcript levels have been reported^16,71^. This indicates a discordance between the p16 transcript and protein expression. In line with this, our data set showed no significant difference in p16 transcript levels between p16-positive and p16-negative regions. Yet, we observed clear differences at the p16 protein level. Importantly, p16 is not a universal marker of senescence. Its abundance can vary across cell types, may be absent in p21 or p53-driven senescence, and can also be induced in non-senescent situations such as tissue stress or neoplasia^43,72,73^. For these reasons, our study does not assume that p16 captures all senescent ovarian cells. Instead, our study identifies a biologically relevant subset of senescent-like populations that increase with age and exhibit an altered ovarian microenvironment. By integrating p16 immunohistochemistry with multiplexed imaging, spatial transcriptomics, and four independent p16-associated gene signatures, we provide a multidimensional framework to identify and map senescent niches in the human ovary.

A key finding from our research is the identification of a strong 32-gene ovarian senotype expression program enriched in p16-positive regions, referred to as BuckSenOvary (*Signature 2*) (Fig. 4d and 7). This senotype includes traditional senescence-associated secretory phenotype (SASP) factors along with extracellular matrix (ECM) modulators. Notably, p16-positive ovarian tissue regions displayed high levels of SERPINE1 (PAI-1), TIMP3, CCN1 (Cyr61), and complement components (CFD, C3), as well as PCOLCE, among others^23,59^. Many of these are well-known mediators of senescence or fibrosis in aging: SERPINE1 and TIMP3 are SASP factors that promote matrix buildup by blocking proteases. CCN1 is a matricellular protein that influences fibrotic remodeling and can trigger fibroblast senescence as a negative feedback mechanism during wound repair. Complement proteins like CFD and C3 are increasingly seen as SASP components that exacerbate inflammation in aging tissues^23,57^. On the other hand, our downregulated *Signature 1* (Fig. 4b and Extended Data Fig. 8a) showed a clear loss of cell-cycle drivers (CCNB1, CCN5, TGFBR3), ribosomal proteins (RPL/RPS family members, EIF4B), and metabolic regulators (SLC7A2, RBP1, PDK4) in p16-positive regions. This finding aligns with the cell-cycle halt and reduced protein synthesis typical of senescent cells^54–56^. Overall, this transcriptional profile combines growth arrest with secretory activation, highlighting key traits of fibroblastic senescence and, for the first time, showcasing them in situ within ovarian tissue. By examining these networks in detail, our data builds on earlier transcriptomic studies of ovarian aging and identifies the specific regions where senescent cells play a role in tissue remodeling. Moreover, the superior performance of BuckSenOvary compared with bone-derived signatures such as SenMayo underscores the importance of organ-specific senotypes for capturing tissue-resident senescence programs.

It is well established in mammalian models, and increasingly supported in humans, that ovarian fibrosis and tissue stiffening increase with age. This rise is driven by changes in the ECM and matrisome^11,66,74–76^. Our findings offer a clear explanation for these long-standing observations. We identified distinct p16-positive senescent areas that serve as focal microenvironments. These areas remodel their surrounding matrix and likely promote fibro-inflammation by secreting a pro-fibrotic senescence-associated secretory phenotype (SASP). The p16-positive regions showed significant fibrotic remodeling. This was not due to more collagen, as postmenopausal ovaries already have high collagen levels. Instead, it resulted from qualitative changes in collagen fiber structure and organization, aligning with stromal stiffening. These architectural changes were sensitively captured by AI-based fibrosis scoring (FibroNest) and were consistent with matrisome enrichment patterns observed in our spatial transcriptomic analyses. We noted an increase in key fibrosis-related genes, including the profibrotic growth factor TGFB1. This factor is a key regulator of collagen organization in various tissues. Along with TGFB1, we observed elevated levels of SASP-related and ECM-modifying factors such as SERPINE1 (PAI-1), PCOLCE, and TIMP3. Additionally, the presence of CD68positive macrophages in p16-positive regions indicates an inflammatory-fibrotic loop that promotes ongoing remodeling. Together, these results show how an increase in fibrosis drivers within senescent niches may contribute to age-related stromal stiffening in the human ovary.

Our data also raise broader questions about the identity and roles of these senescent cells in ovarian aging and disease. The p16-positive niches in postmenopausal ovaries are largely made up of stromal, vascular, macrophage-like, and myofibroblast-like populations, but their origins remain unclear and likely reflect cumulative ovulatory injury, ischemia-reperfusion, inflammation, and hormonal transitions. In other tissues, senescent cells can be acutely protective, limiting proliferation, promoting wound repair, or acting as a barrier to malignant transformation, and have been proposed to act as an anti-cancer failsafe, yet become harmful when they persist and continue to signal through chronic SASP^77–80^. A similar duality may apply in the ovary, particularly where extra-ovarian epithelia are acquired, such as endometriotic implants, cortical inclusion cysts lined by fallopian tube-like epithelium, or other metaplastic foci^81–84^. Senescence at these interfaces could act as a brake on aberrant proliferation or metaplasia, or conversely, create a chronic SASP-rich niche that could foster fibro-inflammation and influence early neoplastic evolution. Given that epithelial ovarian cancers are most commonly diagnosed after menopause^85–87^, it will be important to determine whether BuckSenOvary-like signatures are present at, or adjacent to, early precursor lesions and how they intersect with known fallopian tube-derived pathways of high-grade serous carcinoma.

Several considerations and future directions arise from this work. First, our analysis was performed on a relatively small number of postmenopausal ovaries and is crosssectional, which limits inference about the temporal dynamics of senescence, fibrosis, and ovarian function; longitudinal or peri-menopausal sampling will be important to define how these niches emerge. Second, GeoMx DSP provides ROI-level rather than single-cell resolution, and future studies using higher-resolution spatial platforms (such as CosMx or MERFISH), combined with multiplex imaging, will be needed to resolve the specific p16-positive cell types and their interactions that shape the senescent niche. Third, p16-based detection does not capture all senescent cells and may include stressed but non-senescent populations, underscoring the need to integrate additional senescence markers, functional assays, and interventional approaches that modulate senotype genes or selectively deplete p16-positive cells to test their causal roles. Finally, given the known discordance between mRNA and protein, global proteomics and mass-spectrometry imaging analyses of the aging ovary will be vital. These analyses will map ECM and matrisome remodeling in greater detail and translate this spatial atlas of senescent niches into actionable targets for restoring tissue homeostasis.

In summary, cellular senescence is a defining feature of the aging human ovary. Rare, spatially discrete p16-positive microenvironments co-localize with macrophages and display a conserved SASP/ECM senotype, marked by suppression of cell-cycle/translation and qualitative collagen reorganization without increased bulk. Spatial transcriptomics and AI-guided pathology identify these fibroinflammatory hotspots with high fidelity (notably via BuckSenOvary (Signature 2)), and transcriptomic signatures derived from these analyses can map p16-positive regions in an unsupervised manner across independent samples. Convergence between native p16-positive niches and our doxorubicin-induced ovarian senescence model further supports the robustness of this ovary-specific senotype. Together, these findings link senescence to ovarian stiffening and remodeling and suggest that targeting senescent stromal niches may represent a strategy to preserve postmenopausal ovarian health and narrow the gap between female healthspan and lifespan.

## Methods and Materials

### Human ovarian tissue acquisition and processing

De-identified human ovarian tissue was obtained from the Northwestern University Reproductive Tissue Library (NU-RTL) under an IRB-approved protocol (STU00215770). Ovaries were obtained from females aged 50 to 84 years (mean 64 ± 8 years) undergoing total hysterectomy or salpingo-oophorectomy for various gynecologic conditions (Supplementary Table 1). Individuals with endometriosis, ovarian neoplasia, BRCA mutations, or a history of breast cancer, radiotherapy, or chemotherapy were excluded. Post-operative pathology classified tissues as benign, pre-malignant, or malignant; however, all samples included in this study were confirmed free of ovarian pre-malignancy or malignancy (Supplementary Table 1). Upon collection, the tissue was divided into coronal sections (3-5 mm thick) such that each section contained an outer cortex and inner medulla (Fig. 1a1). In the absence of significant gross pathology as assessed by a certified gynecologic pathologist, varying sizes of the ovarian cross-sections were designated for research and transported to the laboratory on ice in ORIGIO^®^ Handling^TM^ IVF medium (Cooper Surgical Inc., Trumbull, CT, USA).

Ovarian tissue was processed in ORIGIO^®^ Handling^TM^ IVF medium at room temperature. The 3-5mm sections were divided into smaller tissue pieces containing both the cortex and medulla (Fig. 1a1 and 1a2). For certain participants with complete ovarian sections (Fig. 1a1), the sections were divided equally into 8 tissue pieces (n=4, Participant # 1, 2, 3, and 24) while a variable number of tissue pieces were generated for other participants, depending on the availability of tissue (Supplementary Table 1).

### Tissue fixation, histochemical staining, and imaging

The ovarian tissue pieces were fixed in Modified Davidson’s Fixative (mDF) (Electron Microscopy Sciences, Hatfield, PA, USA), at room temperature for 2 hours and then overnight at 4°C. After overnight fixation, the tissue pieces were washed in and transferred to 70% ethanol and stored at 4°C until further processing. The tissue pieces were then dehydrated in an automated tissue processor (Leica Biosystems, Buffalo Grove, IL, USA), embedded in paraffin, and sectioned (5 µm thickness) with a microtome (Leica Biosystems, Buffalo Grove, IL, USA) (Fig. 1a2).

For each tissue piece from all participants, 10 slides were generated, with two 5 µm sections per slide. The 9th and 10^th^ slides from all tissue pieces were stained for hematoxylin & eosin (H&E). All tissue pieces from the participants with 8 tissue pieces (n=4; Participant# 1, 2, 3, and 24), and one tissue piece with the best histology for the remaining participants (n=41), the 8^th^ slide was stained for the p16^INK4A^ antibody by Immunohistochemistry (IHC) (total n=45 participants), and the 7^th^ slide was stained with Picrosirius Red (PSR) for collagen (Supplementary Table 1).

H&E staining was performed using a Leica Autostainer XL (Leica Biosystems, Buffalo Grove, IL, USA). Tissue sections were then cleared with Xylene (Mercedes Scientific, Lakewood Ranch, FL, USA) in three 5-minute incubations and mounted with Cytoseal XYL (Epredia™ Thermo Fisher Scientific, Waltham, MA, USA). IHC was performed with the p16^INK4A^ antibody (1:300 dilution and 2.67µg/mL concentration) (Cat# ENZ-ABS377-0100, Enzo Life Sciences, Farmingdale, NY, USA) (n=45 participants) using an automated IHC stainer (Leica Biosystems, Buffalo Grove, IL, USA) in collaboration with the Pathology Core Facility at Northwestern University. Antibody optimization was performed on native postmenopausal ovarian tissue containing both ovarian cortex and medulla, along with a positive (Cervical Cancer tissue) (Extended Data Fig. 1a) and negative (Extended Data Fig. 1b) control.

For preliminary histological characterization of p16^INK4A^ neighborhoods, manual IHC was performed on sequential 5µm sections (n=2 participants; (Supplementary Table 1)) using the following antibodies: p16^INK4A^ (same as above), p21^CIP1/WAF1^ (1:100 dilution and 2.29µg/mL concentration) (Cat# M720229-02, Dako, Santa Clara, CA, USA), alpha-smooth muscle actin (a-SMA; 1:1000 dilution and 0.007µg/ml concentration) (Cat# PA0943, Leica Biosystems, Deer Park, IL, USA), and CD68 (1:100 dilution and 0.04mg/ml concentration) (Cat# M087601-2, Agilent, Santa Clara, CA, USA) according to our laboratory’s previously established protocol using heat-induced epitope retrieval and antibody detection with a 3’,3’-diaminobenzidine (DAB) Peroxidase Substrate kit (Vector Laboratories, Burlingame, CA)^13^. The slides were then counterstained with hematoxylin (Mercedes Scientific, Lakewood Ranch, FL, USA), cleared with CitriSolv (Decon Labs, King of Prussia, PA, USA), and mounted with Cytoseal XYL.

For PSR staining, tissue sections were deparaffinized with Xylene for two 5-minute incubations, rehydrated with 100% ethanol for three 1-minute incubations, and washed in distilled water. Slides were incubated in Bouin’s fixative (StatLab, McKinney, TX, USA) for 1 hour, Picrosirius Red stain (StatLab, McKinney, TX, USA) for 2 minutes, and 0.5% Glacial Acetic Acid (Fisher Scientific, Pittsburgh, PA, USA) for two 5-second incubations with continuous agitation. The sections were then dehydrated with 100% ethanol for three 10-second incubations, cleared with Xylene for three 1-minute incubations, and mounted with Cytoseal XYL.

To image entire ovarian tissue sections stained with H&E, IHC, and PSR, scans comprised of a series of individual images were taken across the tissue and automatically stitched using a 20X objective on the EVOS FL Auto Cell Imaging System (ThermoFisher Scientific, Waltham, MA, USA). For tissue sections stained with p16^INK4A^, the sections were visualized from one end to the other on the RebelScope Imaging System (Discover ECHO Inc., San Diego, CA, USA) using a 20X objective with 200% optical zoom to map all p16^INK4A^ staining (clusters, isolated punctate staining, staining around vessels or special structures (Fig. 1d). Images of p16^INK4A^ staining throughout a tissue section, as well as on sequential tissue sections, were taken and mapped using either a 20X or 40X objective with 200% optical zoom (Extended Data Fig. 2a and 2b). For PSR quantification, participants with the best visible distinction between cortex and medulla were selected (n=28), and 1-4 ROIs each for cortex and medulla were taken using the 20X objective (Extended Data Fig.11), such that they covered the entire tissue piece. The ROI images were then converted into an RGB format using Fiji (ImageJ2 version 2.14.0/1.54f, Madison, WI, USA), and the second channel, i.e., green channel, was selected. Based on the thresholds for the two youngest and the two participants, a threshold of 125 was applied across all images to select for collagen staining (in red). The percent positive staining was calculated by determining the area of the positive stain relative to the whole ROI, and the values were averaged to obtain individual percentage positive values for cortex (cortex-only ROIs), medulla (medulla-only ROIs), and the whole tissue section (all ROIs).

For digital image labelling, ovarian tissue pieces stained with p16^INK4A^ were scanned in brightfield with a 20X Plan Apo objective using the NanoZoomer Digital Pathology whole slide scanning system (HT-9600) (Hamamatsu City, Japan) at the University of Washington Histology and Imaging Core and in collaboration with Visiopharm^®^ (Broomfield, CO, USA). The Digital Image Analysis (DIA) platform Visiopharm Integrator System (VIS; Ver. 2023.01.1.13563) (Visiopharm, Hørsholm, Denmark) was used to analyze the IHC. Positive staining was detected by binary thresholding and was assigned a yellow color, while negative staining was assigned a blue color. The percent positive staining was calculated by determining the area of the positive stain label relative to the whole tissue section area. For participants with 8 tissue pieces (n=4), an average of the p16^INK4A^ positive area from all 8 tissue pieces was calculated to represent the p16^INK4A^ positive area for that participant.

### Tissue processing for immunofluorescence multiplexing

Johns Hopkins University received eight slides for each donor (n=4). Based on p16 IHC staining performed at Northwestern University, 13 slides exhibiting substantial p16 expression were selected for multiplexed immunofluorescence analysis. Tissue pieces 1, 2, and 4 from donor 1360, tissue pieces 4, 5, and 7 from donor 1368, tissue pieces 4, 5, 6, and 7 from donor 1369, tissue pieces 3, 5, and 8 from donor 2369 were selected based on the p16 IHC staining. Four-micron paraffin sections were baked at 42 °C for 3 hours and dried overnight at room temperature with a desiccator, dewaxed using xylene, rehydrated with a series of alcohols, and concluded with several times of dipping in water. The tissue slides were transferred to a heat-resistant plastic bowl filled with antigen retrieval solution (Vector Laboratories, H-3300-250) and subjected to 20 minutes of heating in a food steamer (Bella).

### Immunofluorescence staining for immunofluorescence multiplexing

p16 (Roche diagnostics, 705-4793), CD68 (Roche diagnostics, 790-2931), and aSMA (Invitrogen, 14-9760-82) were detected with sequential TSA-based immunofluorescence in the first imaging round. Counterstaining was performed with 0.6 mM Hoechst 33342 in Blocker™ casein solution for 15 minutes. The stained tissue sections were then imaged using an inverted fluorescence microscope (see detailed in the Fluorescence Microscopy section). One drop of TBS-T was added to the tissue region to avoid evaporation while imaging. After imaging, fluorophore inactivation steps were performed to reduce the fluorescence signal to the background level. Tissue sections were placed in a transparent box, which was then filled with the bleaching solution containing 2 M H₂O₂ and 3 mM EDTA in PBS at pH 12.5. The transparent box, holding the tissue slides and bleaching solution, is positioned between two 5000 lux light pads (HSK, 615517997868) for one hour to facilitate fluorophore inactivation. In the second imaging round, 53BP1 (Bethyl laboratory, A700-011) was detected with TSA-based immunofluorescence, followed by staining of HMGB1(Abcam, 195010) and Lamin B1 (Abcam, 194108) with directly conjugated antibodies. Counterstaining was performed with 0.6 mM Hoechst 33342 in Blocker™ casein solution for 15 minutes. The stained tissue sections were then imaged using an inverted fluorescence microscope (see detailed in the Fluorescence Microscopy section). One drop of TBS-T was added to the tissue region to avoid evaporation while imaging.

### Fluorescence microscopy for immunofluorescence multiplexing

Fluorescently labelled tissue sections were imaged with a Hamamatsu Flash 4.0 CMOS camera mounted on an inverted research microscope (Ti-E, Nikon). The microscope is equipped with a motorized stage and motorized excitation and emission filters controlled by NIS-Elements (Nikon). Lumencor SpectraX 6 (Lumencor) was used as the light source. For each sample, a custom grid setup was determined to acquire images covering the entire tissue area using an S Fluor 10x microscope objective with an NA of 0.5 (MRF00100, Nikon). For image stitching, the grid step size is set to contain a 10% overlap between adjacent images. The Perfect Focus System (Nikon) was used to maintain a consistent imaging focal plane across the tissue area. Under this microscopic setup, the pixel size of the acquired images was 0.65 mm. The images acquired in each grid were stitched using a previously described method^88,89^.

### Tissue image registration for immunofluorescence multiplexing

This method relies on nuclear images, using the DAPI channel or its equivalent as a reference. The registration process consists of two main steps: global rigid registration followed by local grid-based deformable registration^90^. Global rigid registration was performed on down-sampled images to enhance computational efficiency, while the local deformable registration was applied to full-resolution images to achieve high spatial accuracy. The deformable registration used a grid step size of 500 pixels. The resulting aligned whole-slide images were exported in OME-TIFF format using the libvips library^91^ and qualitatively assessed in QuPath^92^.

### Cell profiling and analysis of immunolabeled images

Nuclei segmentation was performed on the DAPI-stained channel using the pretrained StarDist model^93^. Cell boundaries were defined by expanding each segmented nucleus outward by 4.5 µm (equivalent to 7 pixels). In cases where expanded boundaries overlapped between adjacent nuclei, the boundary was adjusted to the midpoint between them. Image processing and quantification of cellular morphological features were conducted using a custom MATLAB program^89,94^. Morphological features, including cell and nuclear area, aspect ratio, circularity, and equivalent radius, were quantified based on established methods^88,89,94^. Nuclear intensity features, such as mean and total fluorescence intensity, were measured across all aligned channels following background subtraction. Background images were generated for each channel using a 2D median filter with a 7 × 7-pixel window applied to images down-sampled by a factor of 10, and then rescaled to the original resolution.

### Sample preparation for Digital Spatial Profiling (DSP)

Sample preparation followed the NanoString GeoMx DSP slide-preparation user guide (MAN-10150-05, November 2023 updated version) and Merritt et al. 2020. Formalin-fixed paraffin-embedded (FFPE) tissue sections (5 μm) were mounted on Superfrost Plus slides (Thermo Fisher Scientific) and baked at 60 °C for 2 h. Slides were deparaffinised (3 × 5 min in xylene) and rehydrated through graded ethanol (2 × 5 min in 100 % EtOH, 1 × 5 min in 95 % EtOH), followed by a rinse in phosphatebuffered saline (PBS). Antigen retrieval was carried out in 10 mM Tris/1 mM EDTA, pH 9.0, in a laboratory steamer at 100 °C for 15 min. Sections were permeabilised with proteinase K (0.1 mg ml⁻¹, 15 min, 37 °C) and washed in PBS. Slides were hybridised overnight at 37 °C with 250 µl GeoMx probe mix (25 µl Human Whole-Transcriptome Atlas probes, 12.5 µl custom-probe pool, 200 µl Buffer R, 12.5 µl nuclease-free water; NanoString Technologies) under HybriSlip™ coverslips (Grace Bio-Labs). Coverslips were removed by immersion in 2 × SSC containing 0.1 % (v/v) Tween-20, followed by two stringent washes (25 min each, 50 % formamide in 2 × SSC, 37 °C) and a final rinse in 2 × SSC (5 min). Sections were blocked for 1 h at room temperature (RT) in Buffer R supplemented with 7 % (v/v) donkey serum, stained with fluorescently conjugated antibodies for 60 min at RT in the dark, and washed in 2 × SSC for 5 min. Nuclei were counterstained with Syto 83 (Thermo Fisher Scientific; 10 min, RT), slides were rinsed in PBS, and then loaded onto the GeoMx Digital Spatial Profiler (NanoString Technologies) for region-of-interest (ROI) selection and oligonucleotide collection. The study utilized three key antibodies for immunostaining: an anti-CD31 antibody (clone JC/70A) from ABCAM (catalog AB215912) conjugated to Alexa Fluor 694 and used at a 1:100 dilution; an anti-Transgelin antibody (clone SM22-alpha) from Novus (catalog NBP3-121157) conjugated to Alexa Fluor 594 at 1:100 dilution; and Syto-83 nucleic acid stain from Invitrogen (catalog S11364) conjugated to Cy3, applied at a 1:10,000 dilution. The CD31 antibody is mouse-derived, and the Transgelin antibody is sheep-derived^95^.

### GeoMx DSP data acquisition

Digital Spatial Profiling was performed on an automated GeoMx-NGS platform (NanoString Technologies, MAN-10152, November 2023 revision). FFPE slides prepared as above were scanned under three morphology channels, Cy3, Texas Red, and Cy5, to visualise segmentation markers. ROIs were drawn in GeoMx DSP software v2.0, and photocleaved oligonucleotide tags from each ROI were aspirated into individual wells of a 96-well PCR plate.

### Library preparation and sequencing

Oligonucleotide eluates were dried and resuspended in 10 µl nuclease-free (DEPC-treated) water; 4 µl of each eluate served as template for PCR library construction with the GeoMx SeqCode primer mix (NanoString Technologies, MAN-10153-01). The amplification step simultaneously appended Illumina P5/P7 adapter sequences and dual-indexed sample barcodes. PCR products were pooled in equal volumes and subjected to two rounds of purification with AMPure XP beads (Beckman Coulter; 1.2 × bead-to-DNA ratio each round) before elution in 20 µl 10 mM Tris–HCl, pH 8.5. Pooled libraries were sequenced on an Illumina NextSeq 550 (NextSeq 500/550 Mid-Output v2.5 kit) in paired-end mode (27 bp × 27 bp) following the manufacturer’s instructions.

### Data preprocessing and quality control

FASTQ files from 92 regions were processed with GeoMxNGSPipeline (v2.0.21) to generate DCC files. LabWorksheet files and OME-TIFF images were exported from GeoMx DSP. Downstream analyses were performed in R. Data import and quality control used GeoMxWorkflows (v1.8.0). ROIs were excluded if they failed any of the following criteria: >50% of genes not expressed; total reads ≥1,000; minimum negative count ≥1; area ≥1,000; percent trimmed-and-stitched reads ≥80%; aligned reads ≥75%; or percent saturation ≥50%. Features (genes/segments) with low signal were further removed based on the negative-probe distribution and gene detection rate. Counts were then normalized using TMM, and batch effects were corrected with standR (v1.6.0).

### Differential Expression (DE) Analysis of Spatial Transcriptomics

Ninety-two regions of interest (ROIs) were grouped by p16 expression status (p16positive or p16-negative), ovarian region (cortex or medulla), and tissue structure type (stroma, vessels, ovarian surface epithelium (OSE), cyst). Gene expression data were quantile-normalized and log_2_-transformed counts per million (CPM). Differential expression analyses were performed in edgeR, with ROIs partitioned into two groups for each comparison. For each gene, fold change (FC), t-score, raw P value, and Benjamini–Hochberg false discovery rate (FDR) were calculated. Comparisons were first conducted between all p16-positive and p16-negative ROIs, followed by stratified analyses within each ovarian region and each tissue structure type. Genes were deemed significantly differentially expressed if P < 0.05 and log_2_FC > 0.5.

### Pathway enrichment analysis

Pathway enrichment analyses were performed using the clusterProfiler package. Differentially expressed genes were mapped to Gene Ontology (GO), Kyoto Encyclopedia of Genes and Genomes (KEGG), and Hallmark gene sets obtained from MSigDB. For each gene set, normalized enrichment score (NES), P value, and Benjamini-Hochberg false discovery rate (FDR) were calculated.

### p16 signature marker identification

Marker genes were identified using the FindAllMarkers() function in Seurat, which compares each identity group against all others. Analyses were performed for all p16-positive and p16-negative ROIs, and further stratified by p16 status within the ovarian region (cortex or medulla) and tissue structure type (stroma, vessels, ovarian surface epithelium, cyst).

### Evaluation of p16-associated transcriptomic signatures for spatial mapping

The ability of each p16-associated gene signature to discriminate p16-positive regions was assessed using the UCell package for signature scoring. Statistical significance of score differences between p16-positive and p16-negative ROIs across ovarian regions (cortex and medulla) was evaluated using a t-test (P < 0.05). Signature scores were spatially mapped onto tissue images using SpatialOmicsOverlay. For each signature, performance was quantified by (a) its accuracy in distinguishing p16 status and (b) its discriminatory power when stratified by ovarian region. A publicly available senescence-associated signature (SenMayo) was used as a comparator to evaluate the relative performance of the p16associated signatures in classifying p16-positive versus p16-negative regions.

### Fibronest analysis and quantification

For n=10 ovarian tissue pieces (from different participants) with the strongest p16^INK4A^ signal and best clusters (Participant# 1, 2, 6, 7, 8, 9, 10, 11, 12, and 13). To analyze the collagen in the annotated p16-positive and negative regions on the PSR scans, we used the FibroNest quantitative digital pathology platform. This platform used AI-based pathology to assess 12 characteristics related to collagen quantity and structure, 13 morphometric traits related to collagen fibers, and seven attributes related to fibrosis architecture. Each trait representation was captured using a histogram depicting its statistical distribution across all annotated p16positive and -negative regions for all tissue sections and was refined into ∼ 300 quantitative fibrosis traits (qFTs), accounting for parameters such as mean, variance, skewness, and kurtosis, etc. From this pool of ∼300 qFTs, the FibroNest platform generated automated, robust, and continuous scores for fibrosis phenotypic signatures. Similar to the Ph-CFS, the composite scores for each category of collagen content, fiber morphology, and fibrosis architecture, referred to as the collagen composite score (CCS), morphology composite score (MCS), and architecture composite score (ACS), respectively, were assessed.

## Data Availability

All data needed to evaluate the conclusions in the paper are present in the paper and/or the Supporting Information. The spatial transcriptomic data have been uploaded to the SenNet Consortium database.

## Acknowledgements

This work was supported by the NIH Common Fund Cellular Senescence Network (SenNet) program: U54 AG075932 (PI: Schilling, Melov), and UH3 CA275681 (PI: Pei-Hsun Wu). The authors acknowledge the scientific engagement of the late Dr. Judith Campisi and her passionate support for this project.

## Contributions

B.S., F.E.D., P.H.W., M.A.W., and P.R.D. conceived and designed the study (Conceptualization). E.D.S. and M.E.G.P. oversaw the acquisition of postmenopausal ovarian tissue and pathology assessment (Resources, Investigation). P.R.D., U.T., and H.A. performed histological analyses, including p16, H&E, and PSR staining (Investigation). F.W., M.J.K., and P.H.W. developed the multiplex iCLAP workflow, analyzed imaging data, and prepared draft figures (Methodology, Formal Analysis, Visualization). N.M., T.T., and S.M. performed spatial transcriptomic profiling and sequencing (Investigation, Data Curation). G.Z. and B.S.K. provided biospecimens and prepared histological sections (Resources). N.F.M., K.S., M.A.W., and D.F. curated and managed the spatial transcriptomic data, including annotation, cleaning, and integration, and conducted differential expression, pathway enrichment, and spatial clustering analyses (Data Curation, Formal Analysis). M.A.W. and P.R.D. prepared figures and data visualizations (Visualization). M.A.W. wrote the first draft of the manuscript (Writing-Original Draft). M.A.W., P.R.D., B.S., F.E.D., and P.H.W. critically revised and edited the manuscript (Writing-Review & Editing). M.A.W., B.S., F.E.D., P.H.W., S.M., D.F., and D.W. provided supervision and project oversight (Supervision).

## Ethics declarations

Birgit Schilling is on the Advisory Board of MOBILion Systems. Other authors declare no conflicts of interest.

---

**Extended Data Figure 1: Optimization of immunohistochemistry staining for p16 antibody.** Representative images showing optimization of immunohistochemistry staining for p16 antibody (Enzo ABS377-0100) with the corresponding p16 quantification using digital labeling (expressed as a percentage). **a,** In postmenopausal human ovarian tissue (80 years old) **b,** In positive control (Cervical cancer tissue) **c,** Negative control for IHC staining (no primary antibody; 80 years old). Scale bars: Whole sections (a) 1000 mm, (b) and (c) 200 mm; Magnified images (1-4) for (a), (b), and (c) 60 mm.

**Extended Data Figure 2: Mapping p16-positive clusters in human ovarian tissue sections.** Representative images from a 69-year-old participant showing mapping of p16 staining across. **a,** the tissue section that shows p16 positivity in distinct clusters spread throughout the tissue piece. Each cluster is systematically imaged at high magnification (20X). The corresponding H&E image of the section is shown for reference. **b,** sequential sections of the tissue piece showing extension of the p16 clusters through the depth of the tissue. Each tissue section is 5 µm in thickness. Representative images of clusters 3 and 4 from (a) are shown up to 60 mm away from the first section. Scale bars: Whole sections 500 mm; magnified images 60 mm

**Extended Data Figure 3: Heterogeneity of p16 expression in the postmenopausal human ovary. a**, Heterogeneous distribution of p16 staining in an ovarian section from an 80-year-old participant, subdivided into eight pieces. The top row shows H&E, the middle row shows p16 immunohistochemistry (IHC), and the bottom row shows p16 IHC after digital image labelling. Binary thresholding was applied to identify p16-positive regions (yellow) and p16-negative regions (blue). The percentage of p16-positive area is indicated in the top left of each panel. Scale bar, 500 mm. **b**, Representative p16 IHC and digital image labelling in ovarian tissue pieces from two participants aged 54 and 74 years. The percent positive area is shown in the top-left corner of each panel. Scale bars: whole section, 500 mm; magnified regions, 80 mm.

**Extended Data Figure 4: Characterization of p16-positive cells and their niche. a,** Preliminary immunohistochemical characterization of p16-positive regions in a 57- and 73-year-old postmenopausal ovary. **b,** Validation of p16 immunohistochemistry (IHC) staining performed at Northwestern University (top row) using a different antibody with immunofluorescence (IF) on adjacent slides at Johns Hopkins University (bottom row). Multiplex images highlight p16 expression (green) and nuclei (blue). **c,** Examples of multiplex IF images from p16- (top row) and p16+ (bottom row) cortical regions from two different participants. White boxes indicate regions shown at higher magnification.

**Extended Data Figure 5: Mapping p16-positive clusters for the DSP participant**. **a,** Representative images showing the persistence of a p16 positive cluster (outlined in black) across serial sections of the tissue piece from the participant selected for Digital Spatial Profiling using GeoMx (80-years-old). The tissue piece was sliced into 18 serial sections (each 5µm in thickness), and sections 2, 5, 8, 11, 14, and 17 were stained for p16. Other sections were either allocated for GeoMx DSP or left unstained for future analysis. **b,** Annotation of all p16-positive clusters across the tissue section selected for DSP with GeoMx (80-years-old). Annotations on this section were then transferred to the adjacent GeoMx DSP section to annotate p16+ and p16-ROIs for targeted transcriptomic profiling. Scale bars: Whole sections 200 mm; magnified images 60 mm

**Extended Data Figure 6: Spatial transcriptomic data analysis pipeline.** 1) Raw data is preprocessed using GeoMx NGS Pipeline software to generate DCC files and generate the counts matrix, followed by the quality control steps at the ROI and gene level. 2) The filtered data proceeds to background (negative probe) and TMM normalization and batch correction. 3) Differentially expressed genes (DEGs) are identified and used to identify perturbed pathways. 4) Gene expression is mapped back onto the images.

**Extended Data Figure 7: p16 positive cellular component and molecular function analysis. a,** Cellular component analysis of p16+ versus p16-DEGs from all ROIs. **b,** Molecular function analysis of p16+ versus p16-DEGs from all ROIs.

**Extended Data Figure 8. Derivation of p16-associated transcriptomic signatures to map senescent cells. a,** Heatmap of shared downregulated differentially expressed genes (DEGs) across all p16+ regions of interest (ROIs), including cortex and medulla. **b,** Pathway enrichment analysis of shared downregulated DEGs, signature 1. **c,** Pathway enrichment analysis of shared upregulated DEGs, signature 2. **d,** Partial least squares discriminant analysis (PLSDA) of ROIs based on p16 status alone (p16+ vs p16-). **e,** Dot plot of genes contributing to clustering of p16+ and p16-regions in the PLS-DA. **f,** Pathway enrichment of downregulated DEGs in p16+ regions from PLS-DA plot. **g,** Pathway enrichment of upregulated DEGs in p16+ regions from PLS-DA plot

**Extended Data Figure 9. Evaluation of senescence-associated signatures for spatial mapping of p16-positive regions. a,** Unsupervised spatial mapping of each signature to ovarian section 1 using the SpatialOmicsOverlay package. Enrichment scores are overlaid onto tissue architecture; far-left panels indicate reference p16 status annotation for each ROI (p16-, blue; p16+, red). **b, d,** Violin plots showing UCell-derived enrichment scores (“p16 mapping scores”) for each signature in ROIs annotated as p16-(blue) or p16+ (red). **c, e,** Violin plots showing UCell enrichment scores for each signature stratified by anatomical region (cortex vs medulla) and p16 status. This analysis tests whether signatures discriminate p16+ regions independently of tissue region. Violin plots were analyzed using a two-tailed t-test; P< 0.05 was considered significant.

**Extended Data Figure 10. Validating the precision of different p16 signatures to map p16+ve regions in native tissue.** p16 signatures developed in Fig. 4 and evaluated in Fig. 5 were independently used to map in an unsupervised manner onto tissue from two different participants, a 67-year-old participant (a) and a 71-year-old participant (b). **a, c,** Unsupervised spatial mapping of signatures 2 and 4 to ovarian sections onto tissue sections from two different participants using the SpatialOmicsOverlay package. Enrichment scores are overlaid onto tissue architecture. **b, d,** representative H&E and p16-IHC of each participant used to cross-reference p16 ‘hot-spots’ in the GeoMx images (a, c). Zoomed-in p16-IHC images highlight p16 clusters that are potentially highlighted in the GeoMx images.

**Extended Data Figure 11. Quantification of collagen deposition in postmenopausal ovaries using picrosirius red (PSR) staining.** Tissue pieces from postmenopausal ovaries (50-84 years old; n=28) were stained with Picrosirius Red (PSR) stain for collagen, and the staining was quantified to calculate the percentage of area positive for collagen staining. **a,** Images of the whole tissue piece stained with PSR. **b, c,** Representative images of (b) cortex ROIs and (c) medulla ROIs from the corresponding tissue piece in (a). The right panels in (b) and (c) show the same ROI image after a threshold of 125 was applied to highlight collagen (seen in red vs gray areas depicting absence of collagen). The collagen content was then quantified as a percentage of area positive for collagen (red staining). **d,** Quantification of collagen deposition across age in postmenopausal ovaries. Scale bars: Whole scans: 500 mm; Magnified images: 100 mm

## References

1 Garmany, A. & Terzic, A. Global Healthspan-Lifespan Gaps Among 183 World Health Organization Member States. JAMA Netw Open 7, e2450241 (2024). 10.1001/jamanetworkopen.2024.50241

2 Dattani, S. & Rodés-Guirao, L. Why do women live longer than men?, <https://ourworldindata.org/why-do-women-live-longer-than-men> (2023).

3 Peacock, K., Carlson, K. & Ketvertis, K. M. in StatPearls (2025).

4 Gracia, C. R. & Freeman, E. W. Onset of the Menopause Transition: The Earliest Signs and Symptoms. Obstet Gynecol Clin North Am 45, 585–597 (2018). 10.1016/j.ogc.2018.07.002

5 Monteleone, P., Mascagni, G., Giannini, A., Genazzani, A. R. & Simoncini, T. Symptoms of menopause - global prevalence, physiology and implications. Nat Rev Endocrinol 14, 199–215 (2018). 10.1038/nrendo.2017.180

6 El Khoudary, S. R., et al. Menopause Transition and Cardiovascular Disease Risk: Implications for Timing of Early Prevention: A Scientific Statement From the American Heart Association. Circulation 142, e506–e532 (2020). 10.1161/CIR.0000000000000912

7 Malek, A. M. et al. The association of age at menopause and all-cause and cause-specific mortality by race, postmenopausal hormone use, and smoking status. Prev Med Rep 15, 100955 (2019). 10.1016/j.pmedr.2019.100955

8 Ossewaarde, M. E. et al. Age at menopause, cause-specific mortality and total life expectancy. Epidemiology 16, 556–562 (2005). 10.1097/01.ede.0000165392.35273.d4

9 Broekmans, F. J., Soules, M. R. & Fauser, B. C. Ovarian aging: mechanisms and clinical consequences. Endocr Rev 30, 465–493 (2009). 10.1210/er.2009-0006

10 Wang, X., Wang, L. & Xiang, W. Mechanisms of ovarian aging in women: a review. J Ovarian Res 16, 67 (2023). 10.1186/s13048-023-01151-z

11 Amargant, F. et al. Ovarian stiffness increases with age in the mammalian ovary and depends on collagen and hyaluronan matrices. Aging Cell 19, e13259 (2020). 10.1111/acel.13259

12 Converse, A. et al. Multinucleated giant cells are hallmarks of ovarian aging with unique immune and degradation-associated molecular signatures. PLoS Biol 23, e3003204 (2025). 10.1371/journal.pbio.3003204

13 Machlin, J. H. et al. Fibroinflammatory Signatures Increase with Age in the Human Ovary and Follicular Fluid. Int J Mol Sci 22 (2021). 10.3390/ijms22094902

14 Mu, L. et al. Physiological premature aging of ovarian blood vessels leads to decline in fertility in middle-aged mice. Nat Commun 16, 72 (2025). 10.1038/s41467-024-55509-y

15 Fan, X. et al. Single-cell reconstruction of follicular remodeling in the human adult ovary. Nat Commun 10, 3164 (2019). 10.1038/s41467-019-11036-9

16 Jin, C. et al. Molecular and genetic insights into human ovarian aging from single-nuclei multi-omics analyses. Nat Aging 5, 275–290 (2025). 10.1038/s43587-024-00762-5

17 Jones, A. S. K. et al. Cellular atlas of the human ovary using morphologically guided spatial transcriptomics and single-cell sequencing. Sci Adv 10, eadm7506 (2024). 10.1126/sciadv.adm7506

18 Lengyel, E. et al. A molecular atlas of the human postmenopausal fallopian tube and ovary from single-cell RNA and ATAC sequencing. Cell Rep 41, 111838 (2022). 10.1016/j.celrep.2022.111838

19 Liang, J. et al. Ovarian aging at single-cell resolution: Current paradigms and perspectives. Ageing Res Rev 110, 102807 (2025). 10.1016/j.arr.2025.102807

20 Wagner, M. et al. Single-cell analysis of human ovarian cortex identifies distinct cell populations but no oogonial stem cells. Nat Commun 11, 1147 (2020). 10.1038/s41467-020-14936-3

21 Wu, M. et al. Spatiotemporal transcriptomic changes of human ovarian aging and the regulatory role of FOXP1. Nat Aging 4, 527–545 (2024). 10.1038/s43587-024-00607-1

22 Lopez-Otin, C., Blasco, M. A., Partridge, L., Serrano, M. & Kroemer, G. Hallmarks of aging: An expanding universe. Cell 186, 243–278 (2023). 10.1016/j.cell.2022.11.001

23 Basisty, N. et al. A proteomic atlas of senescence-associated secretomes for aging biomarker development. PLoS Biol 18, e3000599 (2020). 10.1371/journal.pbio.3000599

24 Campisi, J. & d’Adda di Fagagna, F. Cellular senescence: when bad things happen to good cells. Nat Rev Mol Cell Biol 8, 729–740 (2007). 10.1038/nrm2233

25 Coppe, J. P., Desprez, P. Y., Krtolica, A. & Campisi, J. The senescence-associated secretory phenotype: the dark side of tumor suppression. Annu Rev Pathol 5, 99–118 (2010). 10.1146/annurev-pathol-121808102144

26 Coppe, J. P. et al. Senescence-associated secretory phenotypes reveal cell-nonautonomous functions of oncogenic RAS and the p53 tumor suppressor. PLoS Biol 6, 2853–2868 (2008). 10.1371/journal.pbio.0060301

27 Gorgoulis, V. et al. Cellular Senescence: Defining a Path Forward. Cell 179, 813–827 (2019). 10.1016/j.cell.2019.10.005

28 Kumari, R. & Jat, P. Mechanisms of Cellular Senescence: Cell Cycle Arrest and Senescence Associated Secretory Phenotype. Front Cell Dev Biol 9, 645593 (2021). 10.3389/fcell.2021.645593

29 O’Reilly, S., Tsou, P. S. & Varga, J. Senescence and tissue fibrosis: opportunities for therapeutic targeting. Trends Mol Med 30, 1113–1125 (2024). 10.1016/j.molmed.2024.05.012

30 Saito, Y., Yamamoto, S. & Chikenji, T. S. Role of cellular senescence in inflammation and regeneration. Inflamm Regen 44, 28 (2024). 10.1186/s41232-024-00342-5

31 Devrukhkar, P. R. et al. A Comprehensive Multiomics Signature of Doxorubicin-Induced Cellular Senescence in the Postmenopausal Human Ovary. Aging Cell, e70111 (2025). 10.1111/acel.70111

32 Patel, S. K. et al. Exosomes Released from Senescent Cells and Circulatory Exosomes Isolated from Human Plasma Reveal Aging-associated Proteomic and Lipid Signatures. bioRxiv (2025). 10.1101/2024.06.22.600215

33 Wang, B., Han, J., Elisseeff, J. H. & Demaria, M. The senescence-associated secretory phenotype and its physiological and pathological implications. Nat Rev Mol Cell Biol 25, 958–978 (2024). 10.1038/s41580-024-00727-x

34 Dimri, G. P. et al. A biomarker that identifies senescent human cells in culture and in aging skin in vivo. Proc Natl Acad Sci U S A 92, 9363–9367 (1995). 10.1073/pnas.92.20.9363

35 Schafer, M. J. et al. Cellular senescence mediates fibrotic pulmonary disease. Nat Commun 8, 14532 (2017). 10.1038/ncomms14532

36 Ogrodnik, M. et al. Cellular senescence drives age-dependent hepatic steatosis. Nat Commun 8, 15691 (2017). 10.1038/ncomms15691

37 Docherty, M. H., Baird, D. P., Hughes, J. & Ferenbach, D. A. Cellular Senescence and Senotherapies in the Kidney: Current Evidence and Future Directions. Front Pharmacol 11, 755 (2020). 10.3389/fphar.2020.00755

38 Ansere, V. A. et al. Cellular hallmarks of aging emerge in the ovary prior to primordial follicle depletion. Mech Ageing Dev 194, 111425 (2021). 10.1016/j.mad.2020.111425

39 Krishnamurthy, J. et al. Ink4a/Arf expression is a biomarker of aging. J Clin Invest 114, 1299–1307 (2004). 10.1172/JCI22475

40 Gonzalez-Gualda, E., Baker, A. G., Fruk, L. & Munoz-Espin, D. A guide to assessing cellular senescence in vitro and in vivo. FEBS J 288, 56–80 (2021). 10.1111/febs.15570

41 Suryadevara, V. et al. SenNet recommendations for detecting senescent cells in different tissues. Nat Rev Mol Cell Biol 25, 1001–1023 (2024). 10.1038/s41580-024-00738-8

42 Tuttle, C. S. L., Luesken, S. W. M., Waaijer, M. E. C. & Maier, A. B. Senescence in tissue samples of humans with age-related diseases: A systematic review. Ageing Res Rev 68, 101334 (2021). 10.1016/j.arr.2021.101334

43 Safwan-Zaiter, H., Wagner, N. & Wagner, K. D. P16INK4A-More Than a Senescence Marker. Life (Basel) 12 (2022). 10.3390/life12091332

44 Sherr, C. J. & Roberts, J. M. CDK inhibitors: positive and negative regulators of G1-phase progression. Genes Dev 13, 1501–1512 (1999). 10.1101/gad.13.12.1501

45 Zhang, S., Ramsay, E. S. & Mock, B. A. Cdkn2a, the cyclin-dependent kinase inhibitor encoding p16INK4a and p19ARF, is a candidate for the plasmacytoma susceptibility locus, Pctr1. Proc Natl Acad Sci U S A 95, 2429–2434 (1998). 10.1073/pnas.95.5.2429

46 Idda, M. L. et al. Survey of senescent cell markers with age in human tissues. Aging (Albany NY) 12, 4052–4066 (2020). 10.18632/aging.102903

47 Liu, Y. et al. Expression of p16(INK4a) in peripheral blood T-cells is a biomarker of human aging. Aging Cell 8, 439–448 (2009). 10.1111/j.1474-9726.2009.00489.x

48 Saul, D. et al. Distinct senotypes in p16- and p21-positive cells across human and mouse aging tissues. EMBO J (2025). 10.1038/s44318025-00601-2

49 Phillips, V., Kelly, P. & McCluggage, W. G. Increased p16 expression in highgrade serous and undifferentiated carcinoma compared with other morphologic types of ovarian carcinoma. Int J Gynecol Pathol 28, 179–186 (2009). 10.1097/PGP.0b013e318182c2d2

50 O’Neill, C. J. et al. High-grade ovarian serous carcinoma exhibits significantly higher p16 expression than low-grade serous carcinoma and serous borderline tumour. Histopathology 50, 773–779 (2007). 10.1111/j.1365-2559.2007.02682.x

51 Horree, N., Heintz, A. P., Sie-Go, D. M. & van Diest, P. J. p16 is consistently expressed in endometrial tubal metaplasia. Cell Oncol 29, 37–45 (2007). 10.1155/2007/868952

52 Wagner, K. D. & Wagner, N. The Senescence Markers p16INK4A, p14ARF/p19ARF, and p21 in Organ Development and Homeostasis. Cells 11 (2022). 10.3390/cells11121966

53 Hara, E. et al. Regulation of p16CDKN2 expression and its implications for cell immortalization and senescence. Mol Cell Biol 16, 859–867 (1996). 10.1128/MCB.16.3.859

54 Payea, M. J., Anerillas, C., Tharakan, R. & Gorospe, M. Translational Control during Cellular Senescence. Mol Cell Biol 41 (2021). 10.1128/MCB.00512-20

55 Lessard, F. et al. Senescence-associated ribosome biogenesis defects contributes to cell cycle arrest through the Rb pathway. Nat Cell Biol 20, 789–799 (2018). 10.1038/s41556-018-0127-y

56 Nishimura, K. et al. Perturbation of ribosome biogenesis drives cells into senescence through 5S RNP-mediated p53 activation. Cell Rep 10, 1310–1323 (2015). 10.1016/j.celrep.2015.01.055

57 Saul, D. et al. A new gene set identifies senescent cells and predicts senescence-associated pathways across tissues. Nat Commun 13, 4827 (2022). 10.1038/s41467-022-32552-1

58 Devrukhkar, P. R. et al. A comprehensive multi-omics signature of doxorubicin-induced cellular senescence in the postmenopausal human ovary. bioRxiv, 2024.2010.2002.616143 (2024). 10.1101/2024.10.02.616143

59 Naba, A. et al. The matrisome: in silico definition and in vivo characterization by proteomics of normal and tumor extracellular matrices. Mol Cell Proteomics 11, M111 014647 (2012). 10.1074/mcp.M111.014647

60 Muss, H. B. et al. p16 a biomarker of aging and tolerance for cancer therapy. Transl Cancer Res 9, 5732–5742 (2020). 10.21037/tcr.2020.03.39

61 McConnell, B. B., Gregory, F. J., Stott, F. J., Hara, E. & Peters, G. Induced expression of p16(INK4a) inhibits both CDK4- and CDK2-associated kinase activity by reassortment of cyclin-CDK-inhibitor complexes. Mol Cell Biol 19, 1981–1989 (1999). 10.1128/MCB.19.3.1981

62 Mitra, J. et al. Induction of p21(WAF1/CIP1) and inhibition of Cdk2 mediated by the tumor suppressor p16(INK4a). Mol Cell Biol 19, 3916–3928 (1999). 10.1128/MCB.19.5.3916

63 Serrano, M., Hannon, G. J. & Beach, D. A new regulatory motif in cell-cycle control causing specific inhibition of cyclin D/CDK4. Nature 366, 704–707 (1993). 10.1038/366704a0

64 Burd, C. E. et al. Monitoring tumorigenesis and senescence in vivo with a p16(INK4a)-luciferase model. Cell 152, 340–351 (2013). 10.1016/j.cell.2012.12.010

65 Sorokina, A. G. et al. Correlations between biomarkers of senescent cell accumulation at the systemic, tissue and cellular levels in elderly patients. Exp Gerontol 177, 112176 (2023). 10.1016/j.exger.2023.112176

66 Maruyama, N. et al. Accumulation of senescent cells in the stroma of aged mouse ovary. J Reprod Dev 69, 328–336 (2023). 10.1262/jrd.2023-021

67 Baker, D. J. et al. Naturally occurring p16(Ink4a)-positive cells shorten healthy lifespan. Nature 530, 184–189 (2016). 10.1038/nature16932

68 Baker, D. J. et al. Clearance of p16Ink4a-positive senescent cells delays ageing-associated disorders. Nature 479, 232–236 (2011). 10.1038/nature10600

69 Bussian, T. J. et al. Clearance of senescent glial cells prevents tau-dependent pathology and cognitive decline. Nature 562, 578–582 (2018). 10.1038/s41586-018-0543-y

70 Yan, H. et al. Primary oocytes with cellular senescence features are involved in ovarian aging in mice. Sci Rep 14, 13606 (2024). 10.1038/s41598-024-64441-6

71 Zhou, C. et al. Single-Cell Atlas of Human Ovaries Reveals The Role Of The Pyroptotic Macrophage in Ovarian Aging. Adv Sci (Weinh) 11, e2305175 (2024). 10.1002/advs.202305175

72 Hall, B. M. et al. p16(Ink4a) and senescence-associated beta-galactosidase can be induced in macrophages as part of a reversible response to physiological stimuli. Aging (Albany NY) 9, 1867–1884 (2017). 10.18632/aging.101268

73 Sharpless, N. E. & Sherr, C. J. Forging a signature of in vivo senescence. Nat Rev Cancer 15, 397–408 (2015). 10.1038/nrc3960

74 Dipali, S. S. et al. Proteomic quantification of native and ECM-enriched mouse ovaries reveals an age-dependent fibro-inflammatory signature. Aging (Albany NY) 15, 10821–10855 (2023). 10.18632/aging.205190

75 Shen, L., Liu, J., Luo, A. & Wang, S. The stromal microenvironment and ovarian aging: mechanisms and therapeutic opportunities. J Ovarian Res 16, 237 (2023). 10.1186/s13048-023-01300-4

76 Umehara, T. et al. Female reproductive life span is extended by targeted removal of fibrotic collagen from the mouse ovary. Sci Adv 8, eabn4564 (2022). 10.1126/sciadv.abn4564

77 Calcinotto, A. et al. Cellular Senescence: Aging, Cancer, and Injury. Physiol Rev 99, 1047–1078 (2019). 10.1152/physrev.00020.2018

78 Piskorz, W. M. & Cechowska-Pasko, M. Senescence of Tumor Cells in Anticancer Therapy-Beneficial and Detrimental Effects. Int J Mol Sci 23 (2022). 10.3390/ijms231911082

79 Pulido, T., Velarde, M. C. & Alimirah, F. The senescence-associated secretory phenotype: Fueling a wound that never heals. Mech Ageing Dev 199, 111561 (2021). 10.1016/j.mad.2021.111561

80 Wilkinson, H. N. & Hardman, M. J. Cellular Senescence in Acute and Chronic Wound Repair. Cold Spring Harb Perspect Biol 14 (2022). 10.1101/cshperspect.a041221

81 Cao, L. et al. The controversial role of senescence-associated secretory phenotype (SASP) in cancer therapy. Mol Cancer 24, 283 (2025). 10.1186/s12943-025-02475-8

82 Dubeau, L. The cell of origin of ovarian epithelial tumours. Lancet Oncol 9, 1191–1197 (2008). 10.1016/S1470-2045(08)70308-5

83 Pavone, M. E. & Lyttle, B. M. Endometriosis and ovarian cancer: links, risks, and challenges faced. Int J Womens Health 7, 663–672 (2015). 10.2147/IJWH.S66824

84 Zhang, S. et al. Both fallopian tube and ovarian surface epithelium are cellsof-origin for high-grade serous ovarian carcinoma. Nat Commun 10, 5367 (2019). 10.1038/s41467-019-13116-2

85 Ali, A. T., Al-Ani, O. & Al-Ani, F. Epidemiology and risk factors for ovarian cancer. Prz Menopauzalny 22, 93–104 (2023). 10.5114/pm.2023.128661

86 Arora, T., Mullangi, S., Vadakekut, E. S. & Lekkala, M. R. in StatPearls (StatPearls Publishing Copyright © 2025, StatPearls Publishing LLC., 2025).

87 Rampersad, A. C., Wang, Y., Smith, E. R. & Xu, X.-X.

88 Wu, P. H. et al. Single-cell morphology encodes metastatic potential. Sci Adv 6, eaaw6938 (2020). 10.1126/sciadv.aaw6938

89 Wu, P. H. et al. Evolution of cellular morpho-phenotypes in cancer metastasis. Sci Rep 5, 18437 (2015). 10.1038/srep18437

90 Kiemen, A. L. et al. CODA: quantitative 3D reconstruction of large tissues at cellular resolution. Nat Methods 19, 1490–1499 (2022). 10.1038/s41592-022-01650-9

91 Martinez, K. & Cupitt, J. in *IEEE International Conference on Image Processing* 2005 Vol. 2 II-574 (2005).

92 Bankhead, P. et al. QuPath: Open source software for digital pathology image analysis. Sci Rep 7, 16878 (2017). 10.1038/s41598-017-17204-5

93 Schmidt, U., Weigert, M., Broaddus, C. & Myers, G. in Medical Image Computing and Computer Assisted Intervention. (2018).

94 Phillip, J. M. et al. Biophysical and biomolecular determination of cellular age in humans. Nat Biomed Eng 1 (2017). 10.1038/s41551-017-0093

95 Merritt, C. R. et al. Multiplex digital spatial profiling of proteins and RNA in fixed tissue. Nat Biotechnol 38, 586–599 (2020). 10.1038/s41587-020-0472-9

