## Supplementary material for "Senescence-Linked Fibrosis in the Aging Human Ovary Revealed by p16-Based Histological Profiling and Spatial Transcriptomics": Suppl files 1-11

**Extended Data Figure 1.** Optimization of immunohistochemistry staining for p16 (For Figure 1)

**a)** Human ovarian tissue piece

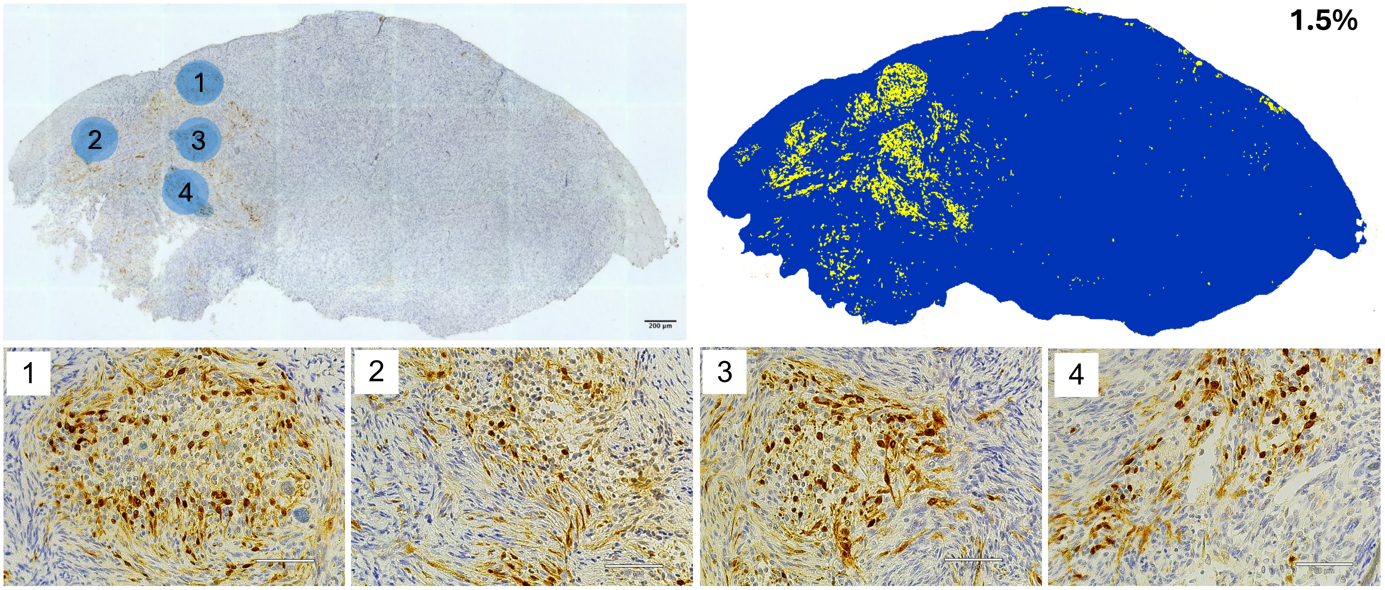


**b)** Positive Control (Cervical Cancer)

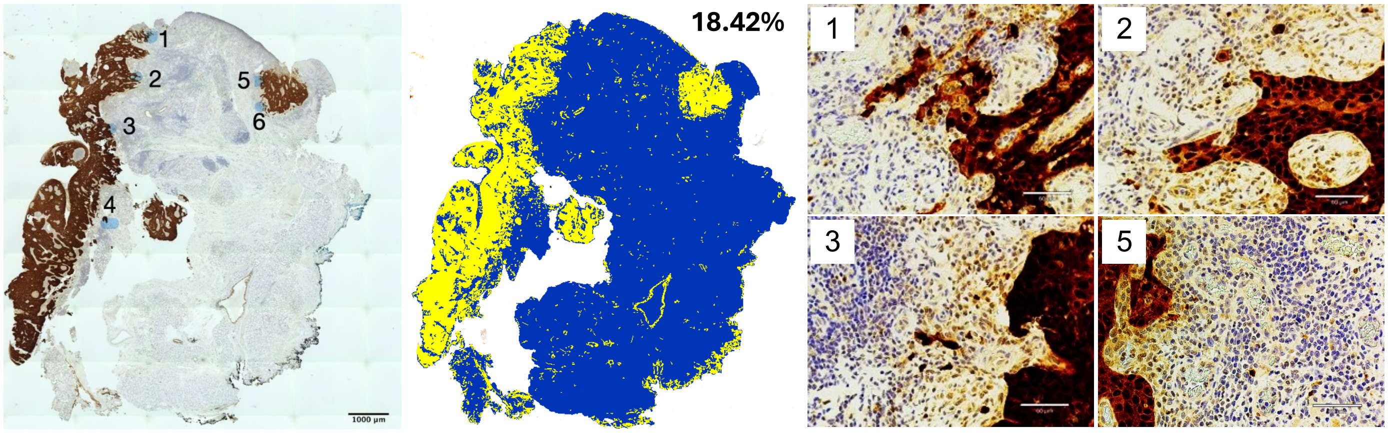


**c)** Negative Control (Human ovarian tissue piece; no primary antibody)

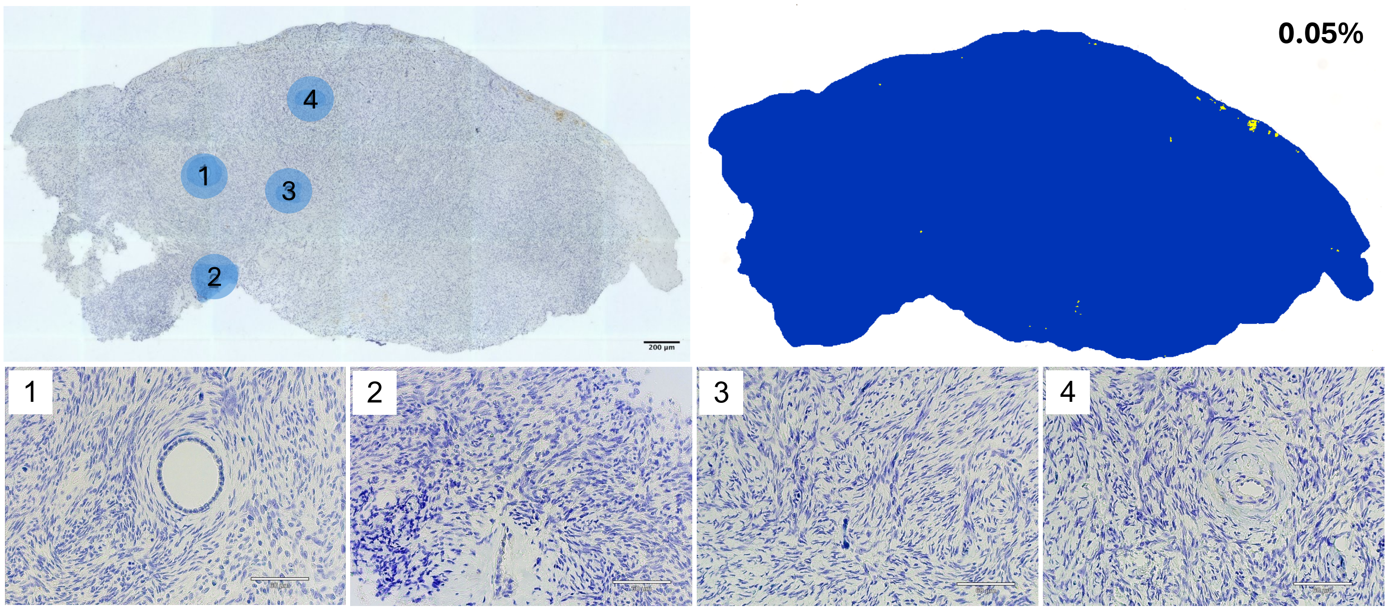

**Extended Data Figure 2:** Mapping p16 positive clusters in human ovarian tissue pieces (For Figure 1)

**a)** Mapping all p16 positive clusters across a human ovarian tissue piece

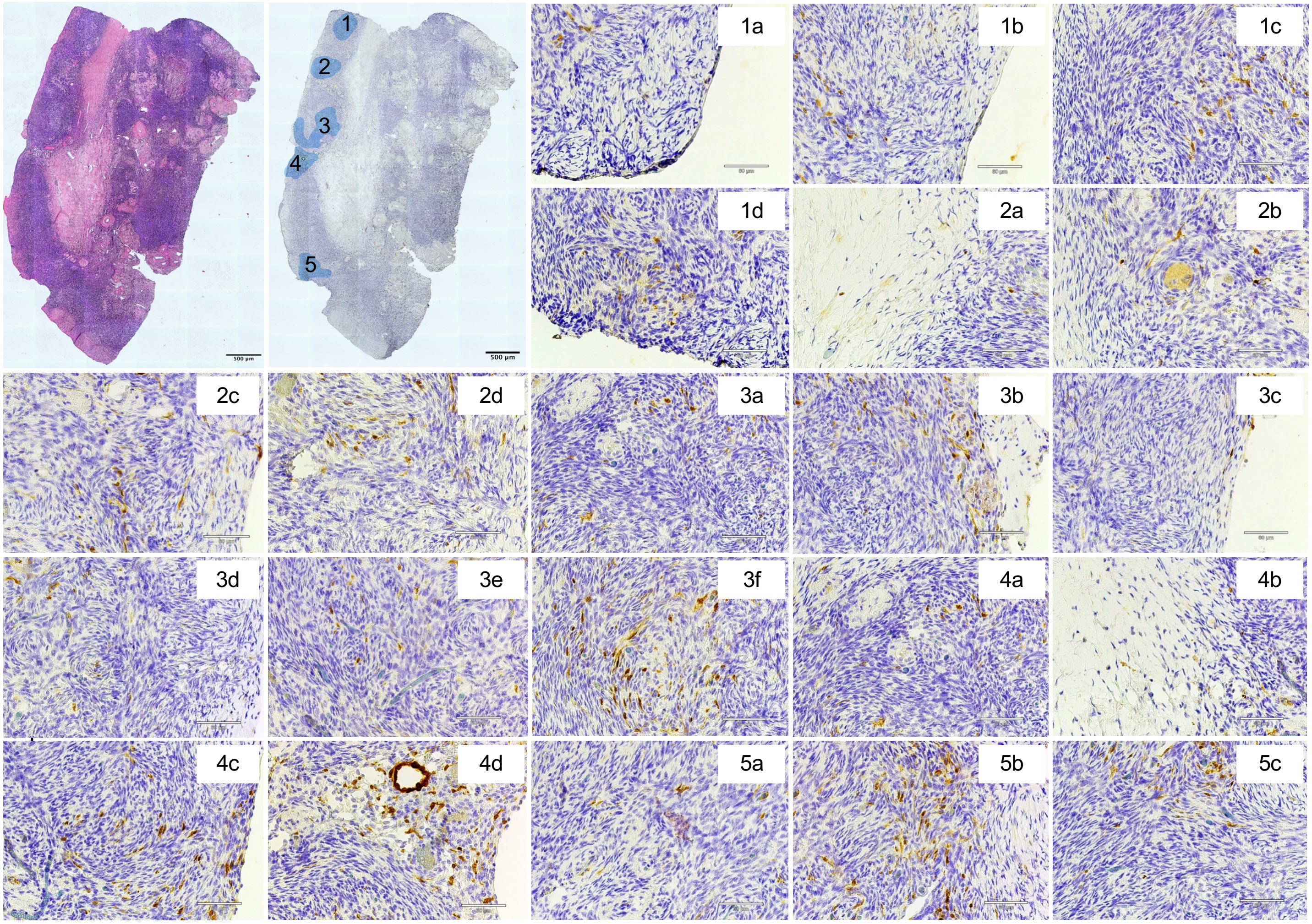


**b)** Mapping p16 positive clusters across sequential sections (Clusters 3 and 4)

+15 µm +30 µm +45 µm +60 µm)

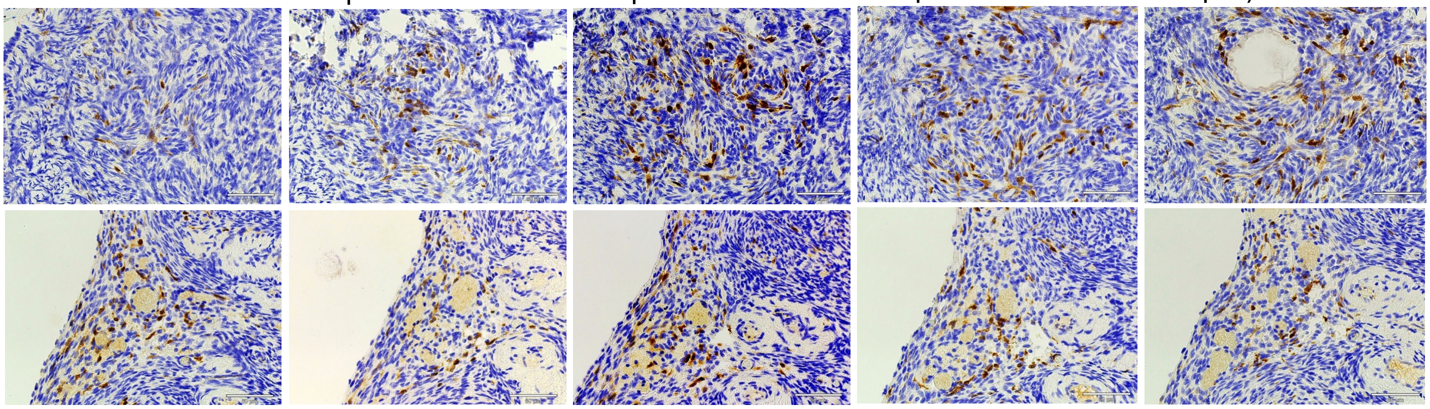

**Extended Data Figure 3:** Heterogeneity of p16 expression in the postmenopausal human ovary

**a)** p16 expression in the postmenopausal human ovary is heterogeneous

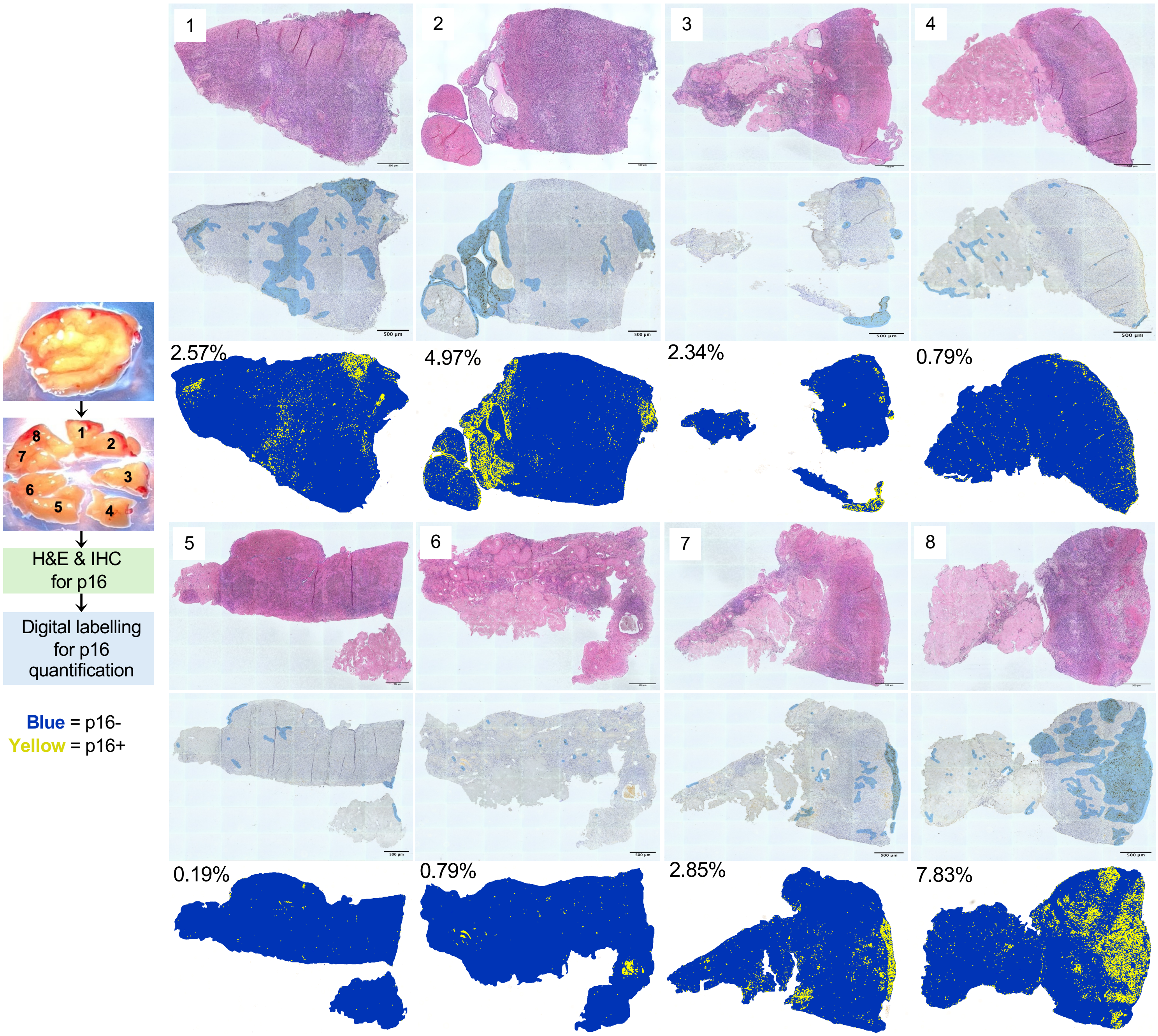


**b)** p16 expression evaluated across different ages

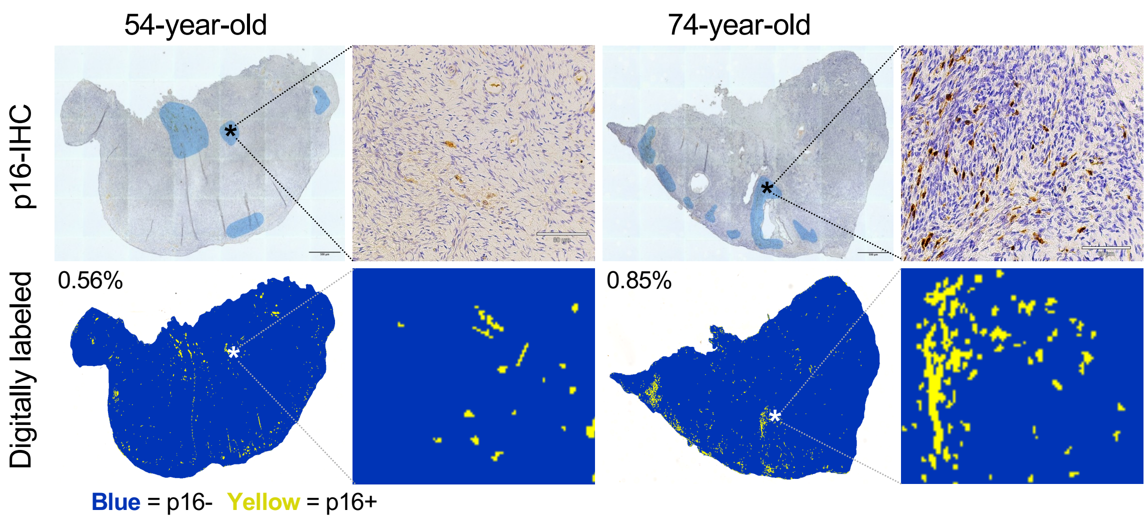

**Extended Data Figure 4:** Characterization of p16-positive cells and their niche

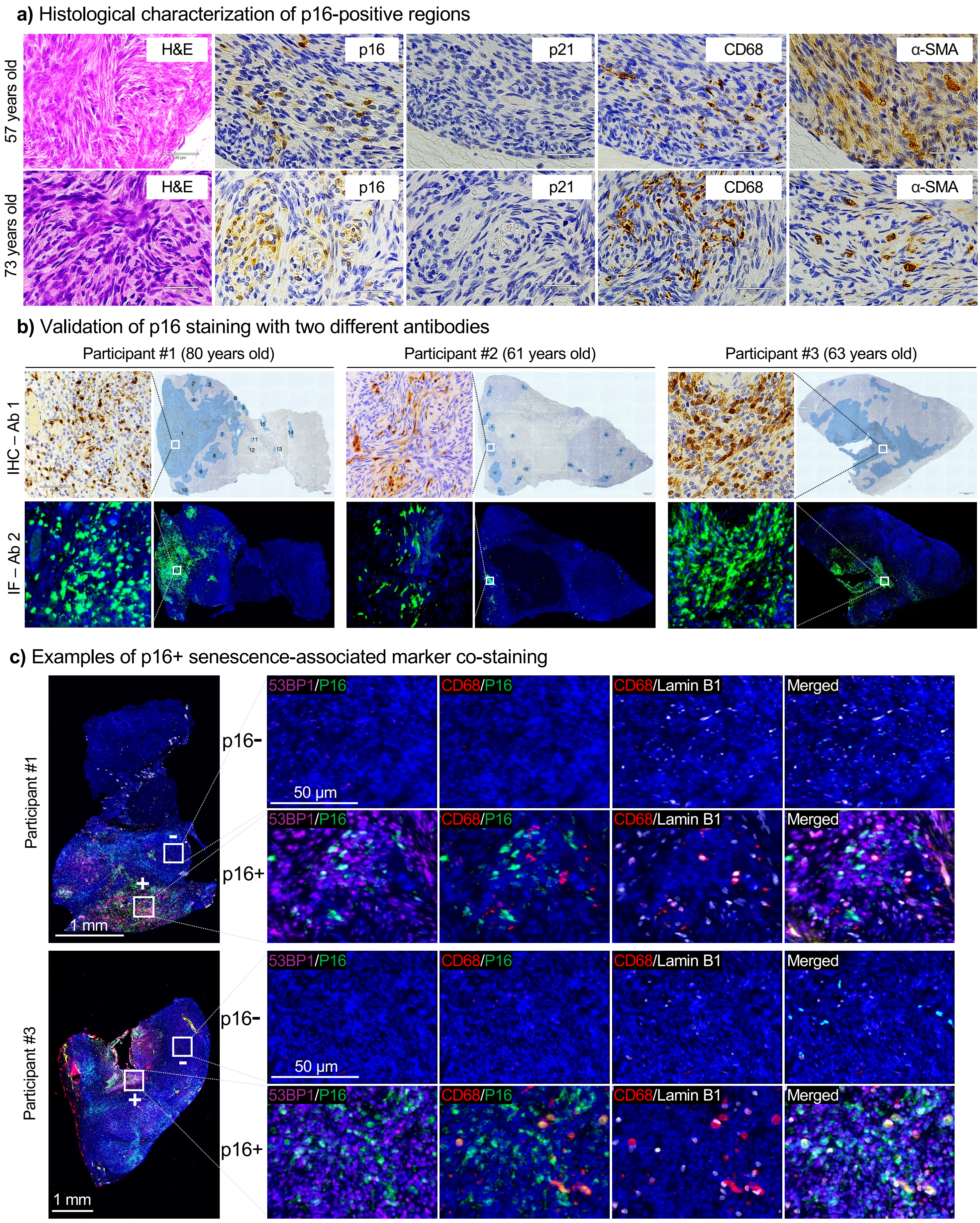

**Extended Data Figure 5:** Mapping p16 positive clusters for the DSP participant

**a)** p16+ clusters persist across several sequential tissue sections

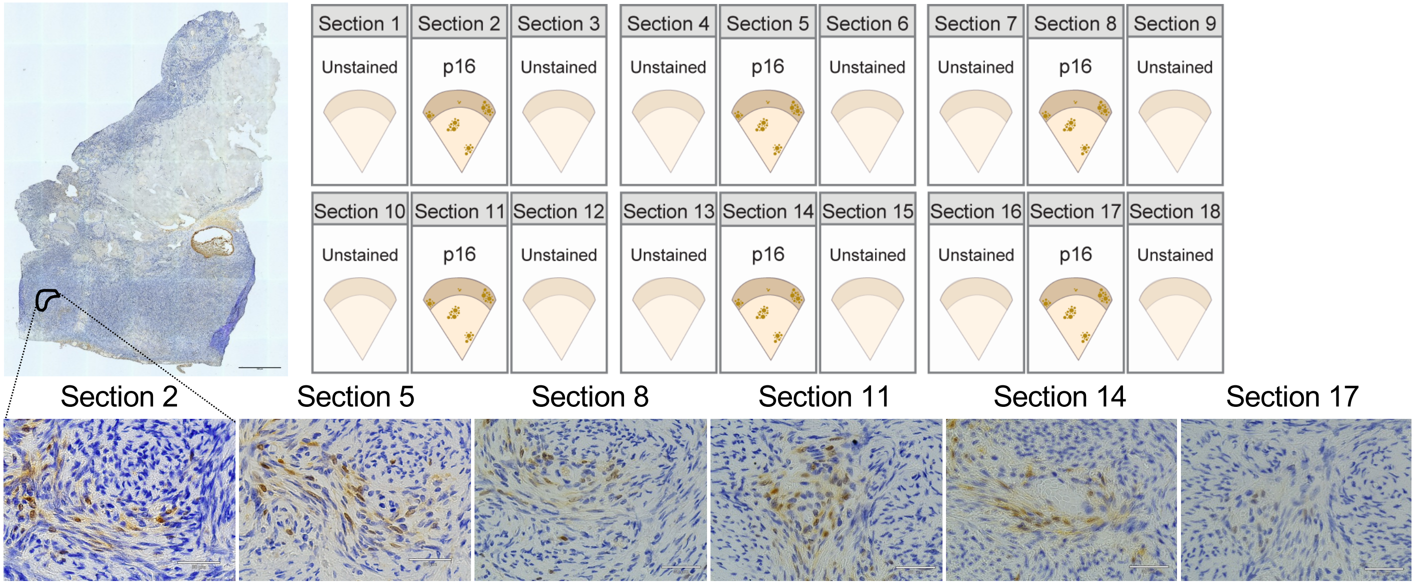


**b)** Annotating p16+ clusters within a tissue section for GeoMx ROI selection

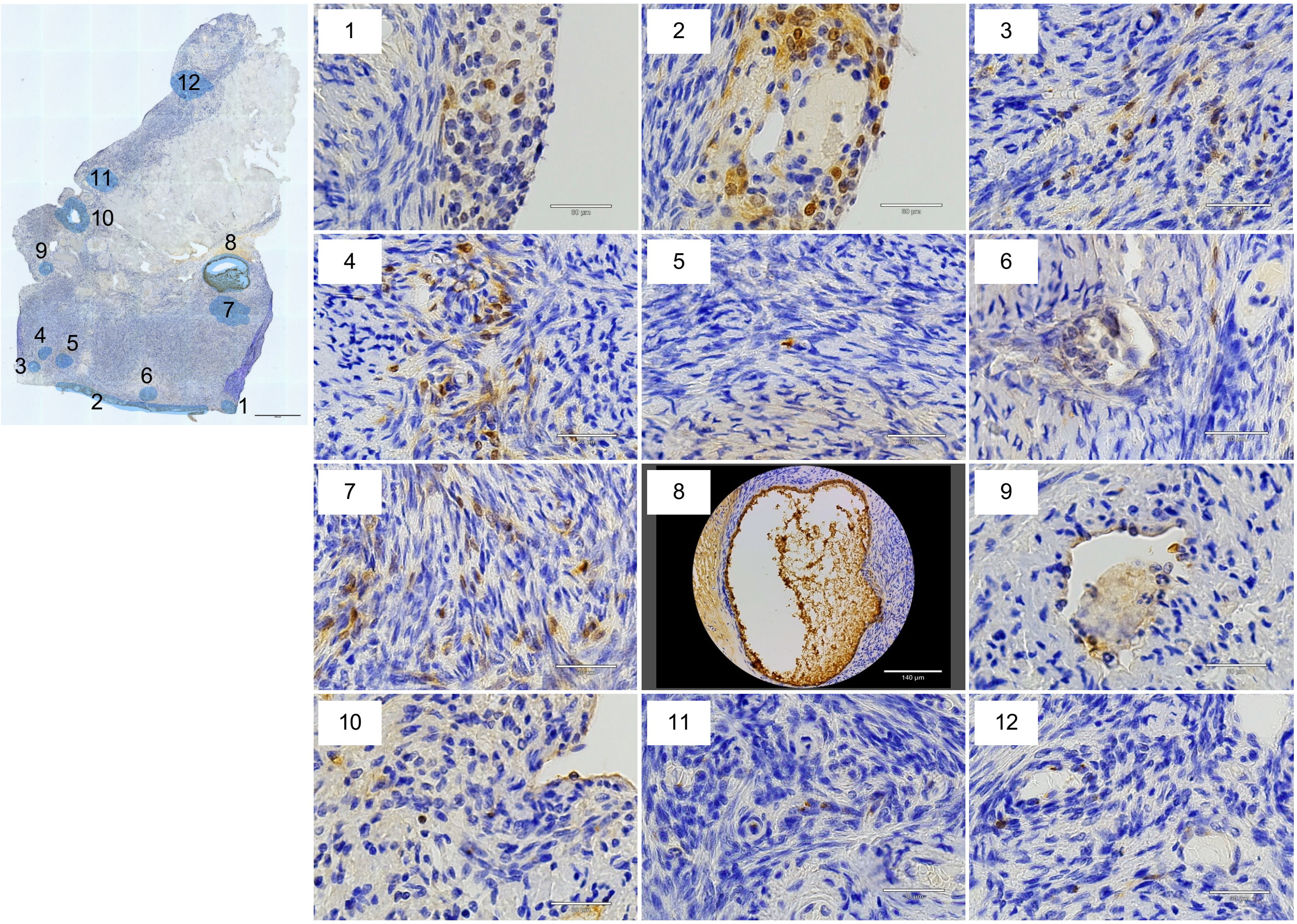

**Extended Data Figure 6:** Spatial transcriptomic data analysis pipeline

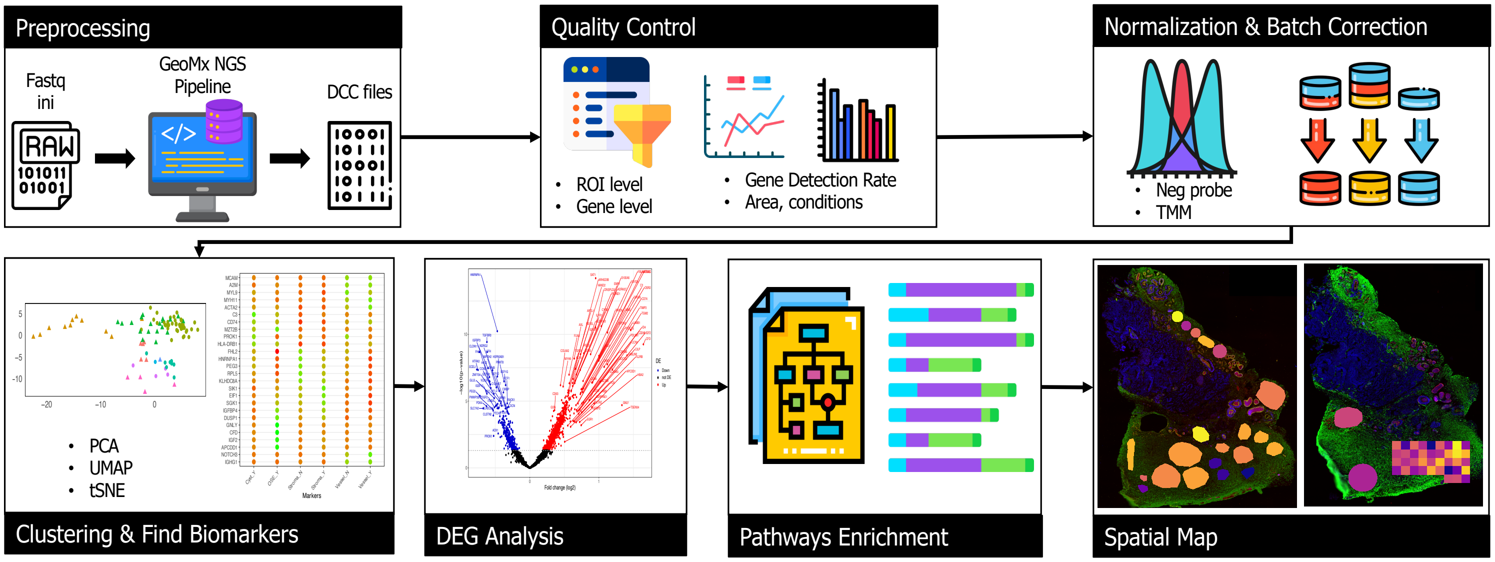

**Extended Data Figure 7:** p16 positive cellular component and molecular function analysis

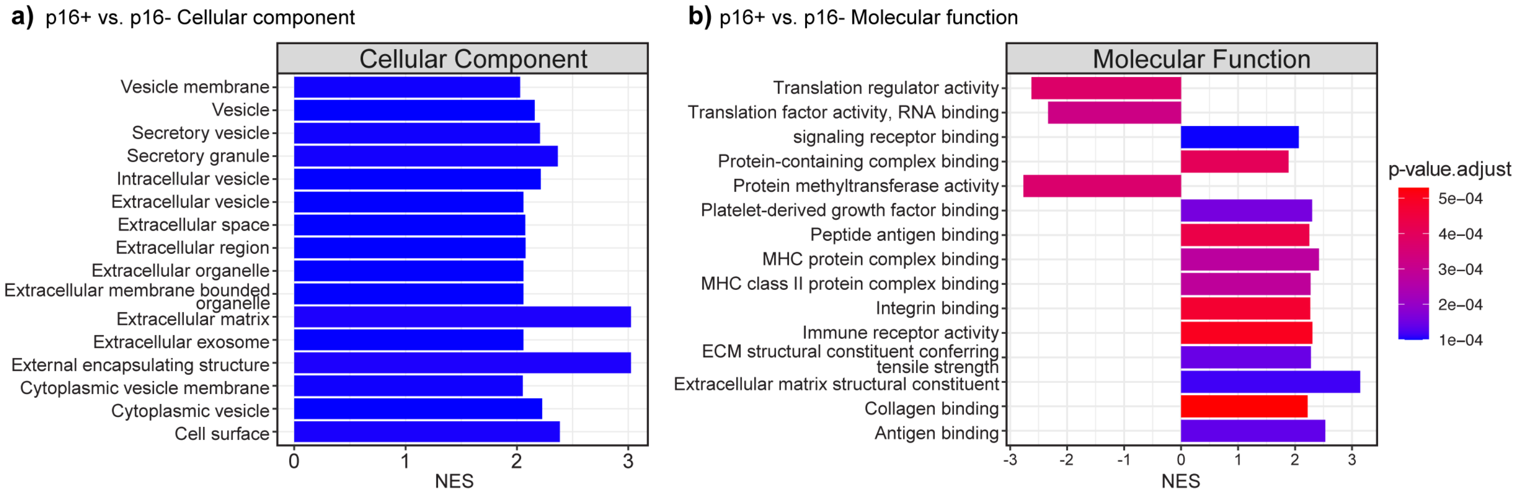

**Extended Data Figure 8:** Derivation of p16-associated transcriptomic signatures to map senescent cells

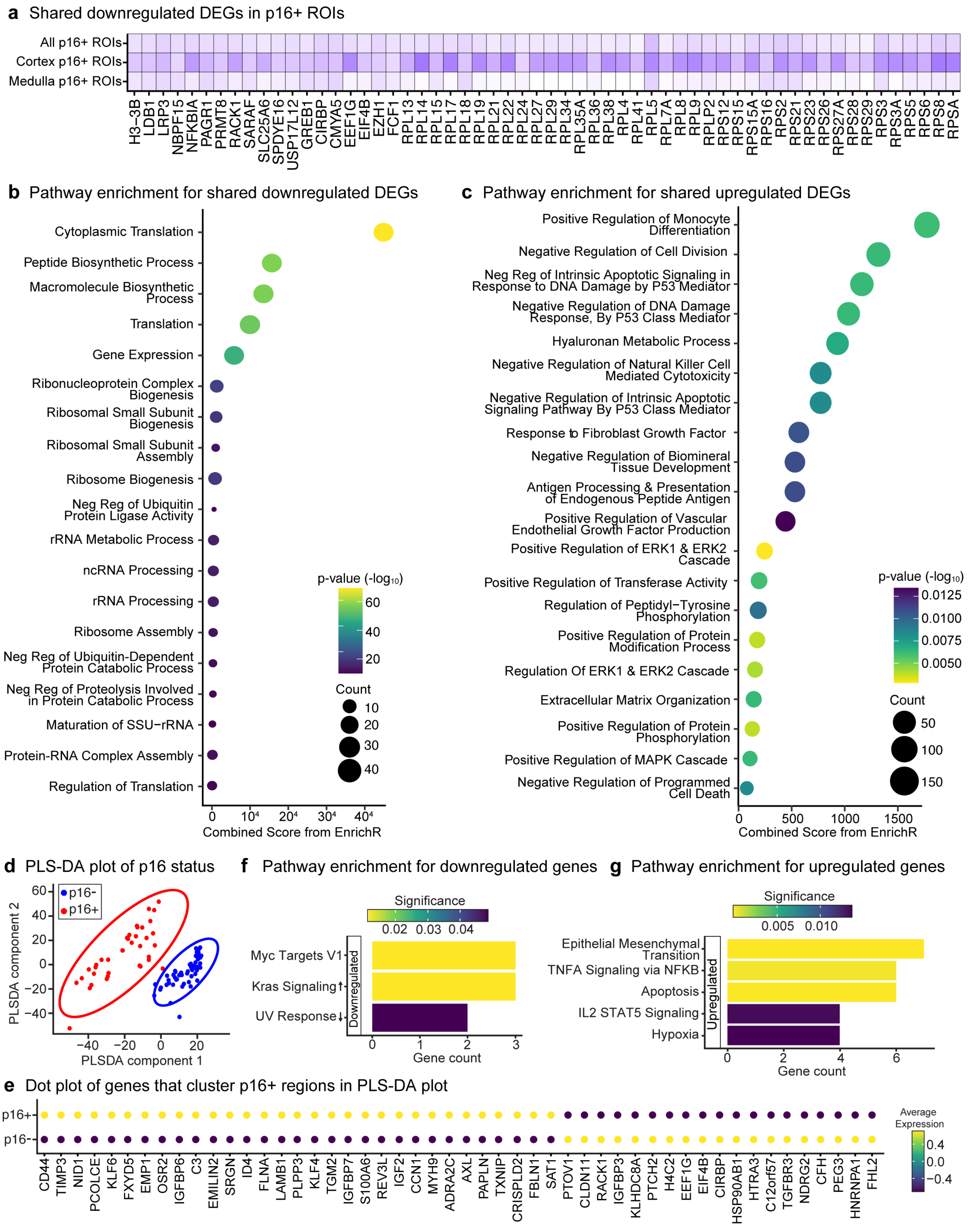

**Extended Data Figure 9:** Evaluation of senescence-associated signatures for spatial mapping of p16 positive regions. (For figure 5)

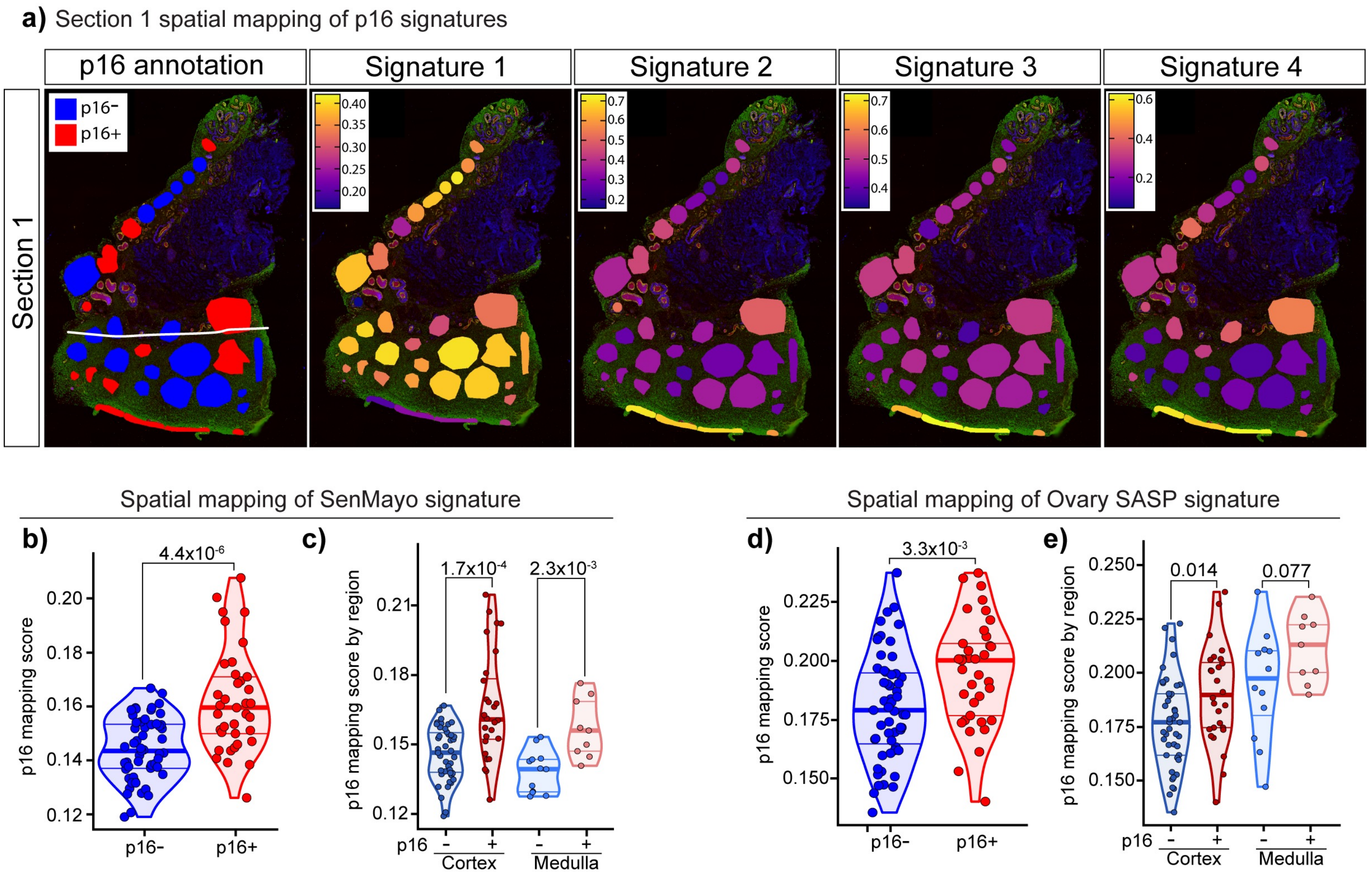

**Extended Data Figure 10:** Validating the precision of different p16 signatures to map p16 positive regions in native tissue (For Figure 5)

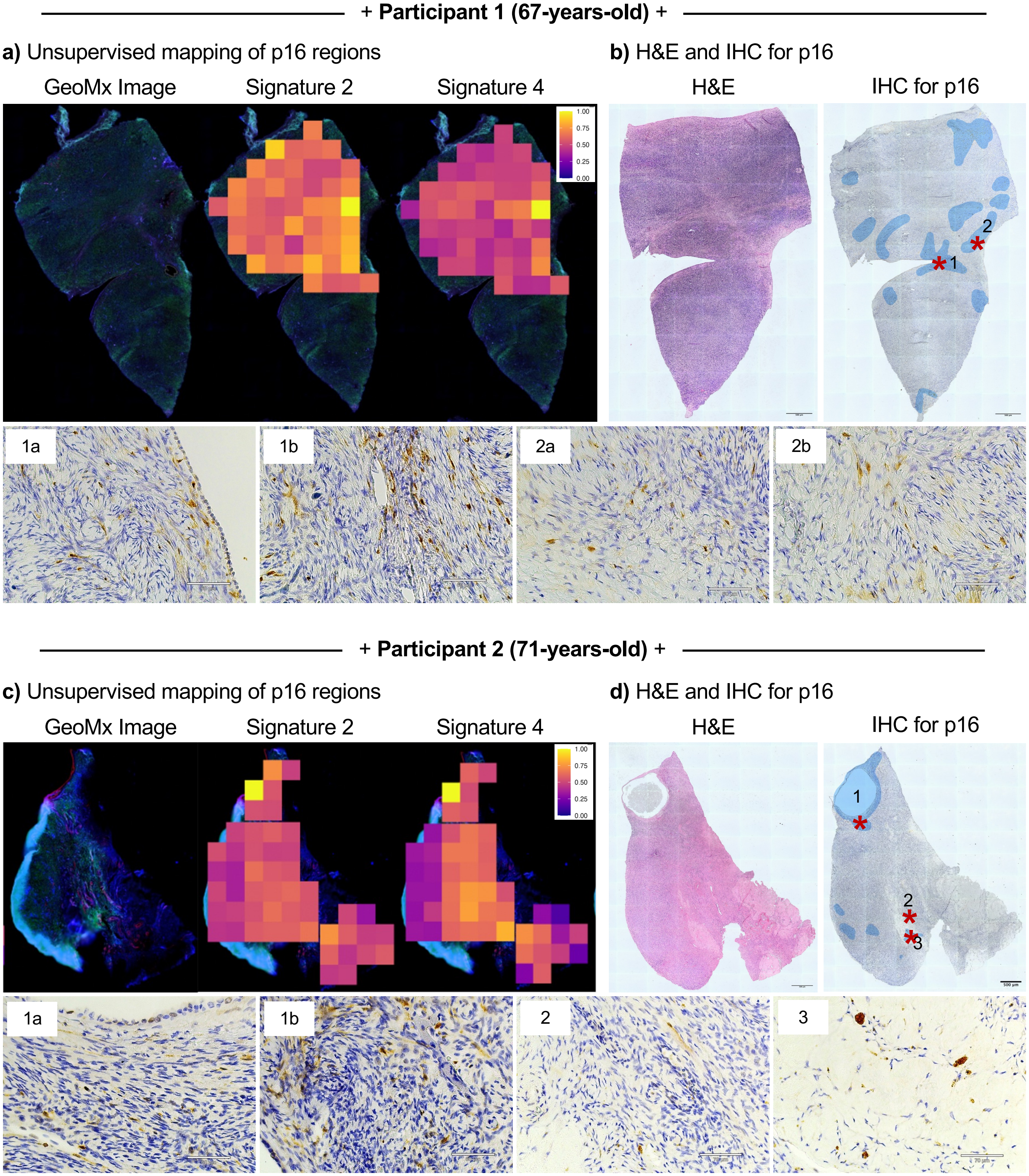

**Extended Data Figure 11:** Histological quantification of collagen deposition in postmenopausal ovaries using picrosirius red (For Figure 6)

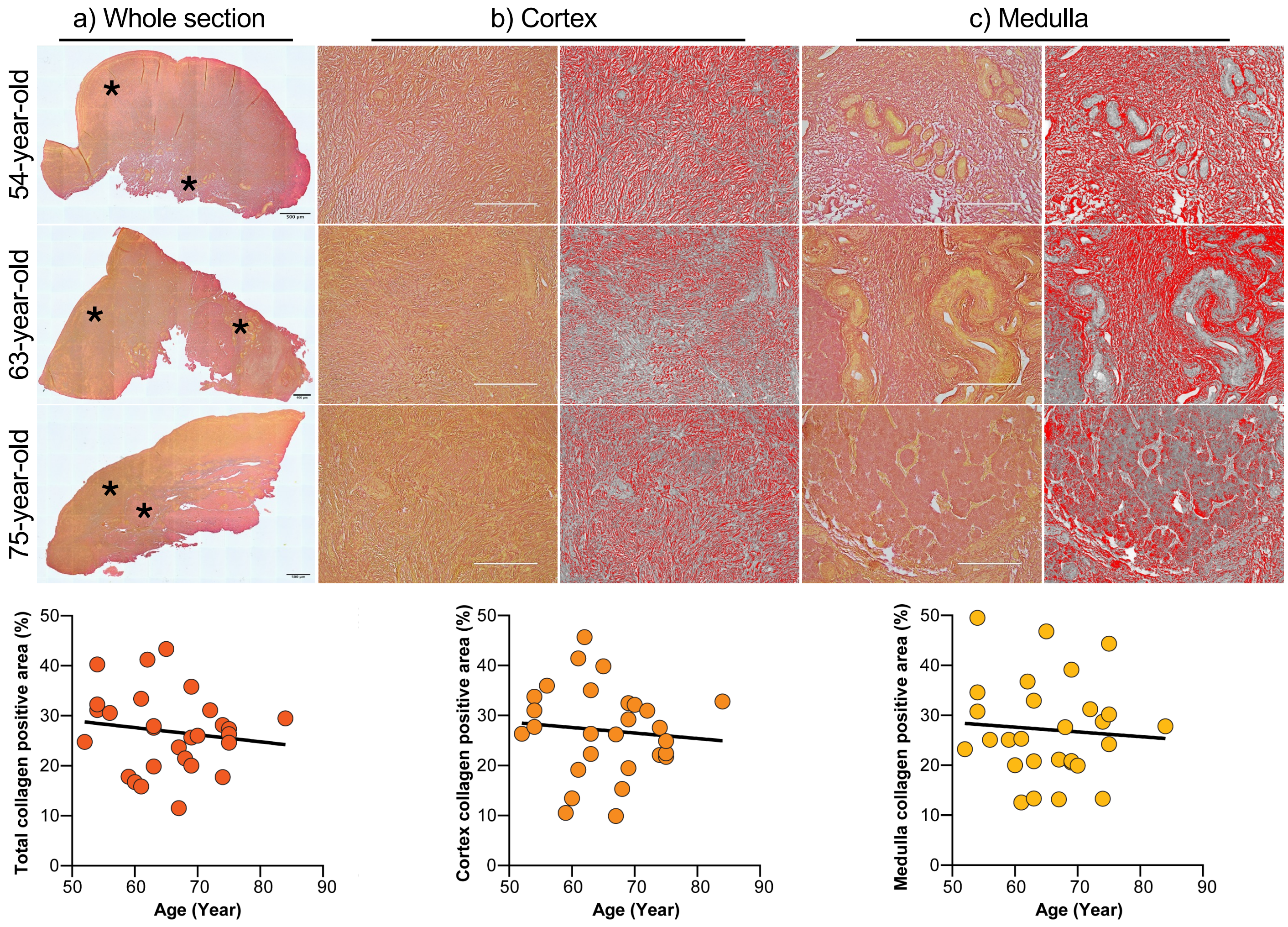
